# Widespread immune systems protect bacteria against conjugative plasmids

**DOI:** 10.64898/2026.07.06.735840

**Authors:** Ludovic Poiré, Julie Baltenneck, Salomé Guillory, Baptiste Clement, Julien Cayron, Christian Lesterlin, François Rousset, Xavier Charpentier

## Abstract

Conjugative plasmids are a class of mobile genetic elements capable of efficient transfer between bacterial cells. Although they can introduce beneficial traits such as antibiotic resistance to recipients, they may also behave as genetic parasites. Bacteria would thus be expected to have evolved barriers to plasmid conjugation. However, the distribution of these barriers and their underlying mechanisms remain poorly understood. Here, we performed a large-scale analysis of 364 diverse strains of the opportunistic pathogen *Acinetobacter baumannii* as recipients of the broad-host-range conjugative plasmids R388 and RP4. Major variations in host susceptibilities to conjugation, with limited phylogenetic association, suggested multiple and fast-evolving plasmid-specific barriers. Functional genetic analyses revealed a role for core genes, pointing to epistasis or genetic background effects. This is illustrated by the previously unrecognized role of H-NS expression in alleviating conjugation barriers in a strain-dependent manner. Most importantly, we identified three novel immune systems protecting bacteria against conjugation by R388 and RP4. Their patchy distribution within the species, and that of their homologs across bacteria, indicate that they are part of a dynamic repertoire of immune systems against conjugation. While the Ishtar system promotes plasmid loss through putative HEPN nuclease domains, Namtar and Attar sense distinct components of the R388 type IV secretion system (T4SS) to trigger a non-proliferative, energetically depleted state, analogously to the abortive infection response of anti-phage defenses. Live imaging of conjugation showed Namtar halting cell division in *Escherichia coli* recipients, conferring population-level immunity against plasmid spread via horizontal and vertical transmission. The existence of immune systems specifically targeting T4SS components suggests that conjugative plasmids impose a selective disadvantage greater than previously thought. This work reveals an additional layer of bacterial immunity directed at a class of genetic elements driving dissemination of antibiotic resistance.

## Introduction

Bacterial genomes harbor multiple mobile genetic elements playing an important role in genome evolution (*1*). Among these, conjugative plasmids are episomal DNA replicating independently of the chromosome (*2*). They encode type IV secretion systems (T4SS) that mediate their horizontal transfer by a process known as conjugation (*3*, *4*). A donor cell forms a mating pair with a recipient, transfers a single-stranded DNA copy of the plasmid via the T4SS to the new host where complementary strand synthesis restores the double-stranded replicon (*5*). Importantly, broad-host range plasmids can transfer within and between species, allowing genes to flow across distant and diverse microbial populations (*6*). As they bring along their cargo genes, conjugative plasmids can provide new traits to their recipients, some of which may be beneficial to their new hosts, such as antibiotic resistance (*7*). Indeed, conjugative plasmids are the primary drivers of resistance gene dissemination among ESKAPE pathogens (*8*, *9*). However, conjugative plasmids may also incur a fitness cost (*10*) or even conflict with their host by interfering with bacterial processes such as motility, natural transformation or apparatus used to kill competitors (*11–13*). Bacteria would thus be expected to have evolved barriers to conjugation (*14*). While the field of bacterial immunity has documented a large repertoire of defenses acting as barriers to phage infection, comparatively little is known about whether have evolved specific barriers to conjugation. For instance, CRISPR-Cas systems and restriction–modification are DNA-targeting prokaryotic immune systems that can cleave plasmid DNA entering by conjugation (*15–17*) but are mostly known for their role in defending against phage infection (*18*, *19*). Other recently discovered anti-plasmid systems also function through generic nucleic acid recognition (*19*). These include Wadjet which can recognize and cleave large circular DNA (*20*, *21*) and DdmDE which eliminates plasmids (*23*) through DNA-guided cleavage using a short prokaryotic argonaute (pAgo) module (*22*, *23*). Other pAgo (*24*) and Lamassu/DdmABC (*25*, *26*) appear as anti-plasmid defenses but can also display anti-phage activity. Beyond defense systems, capsule polysaccharides and other cell-surface structures can limit conjugation by physically impairing mating-pair formation, but may have evolved to confer protection against phage predation or the animal immune system (*27*, *28*). Conjugative plasmids encode surface exclusion mechanisms to interfere with mating pair formation, and entry exclusion mechanisms to block plasmid DNA transfer after productive cell contact, but these adaptations are plasmid-driven strategies meant to reduce the cost of conjugation on their bacterial host (*29*, *30*). Barriers to conjugation and their origin remain poorly documented, limiting our ability to determine whether they represent adaptations to limit plasmid transmission, or incidental by-products of other selective pressures.

In this work, we followed a comprehensive, species-scale approach to investigate barriers to conjugative transfer in the ESKAPE pathogen *Acinetobacter baumannii.* Using a large and diverse panel of strains and high-throughput mating assays, we quantified the variation in conjugation efficiency across the species, revealing the dynamic nature of barriers to conjugation. Genome-wide transposon mutagenesis revealed the genetic determinants of the barriers and identified novel and specific immune systems against conjugative plasmids. This work hence offers new insights into how bacterial immunity can shape the spread of conjugative plasmids.

## Results

### Species-wide determination of *A. baumannii* susceptibility to plasmid acquisition by conjugation reveals fast-evolving barriers

We measured the susceptibility of 364 A. *baumannii* strains (Ab-One collection, **Table S1**) to conjugative transfer of the broad-host-range plasmids R388 (IncW) and RP4 (IncP-1alpha). Both plasmids share a Type T Mating Pair Formation (MPF) type that mediates efficient transfer on solid surfaces. To enable selection of transconjugants, we cloned the *aac*(*4*) resistance gene into each plasmid (RP4* and R388*, hereafter simply noted RP4 and R388); this confers apramycin resistance, a phenotype that is rare in *A. baumannii*. The strains, drawn from multiple environmental and clinical sources, represent a pangenome of 27,080 gene families, covering over 50% of the species’ known genetic diversity (*31*). Phylogenetic reconstruction yielded a tree of mixed topology, containing both well-defined clonal lineages and highly divergent strains (*32*). Each *A. baumannii* strain was mated with an *Escherichia coli* donor carrying either R388 or RP4 and assigned a conjugation score from 0 to 4 (**Fig. 1A**), serving as proxy for transfer frequency (see **Fig. S1** for validation). The conjugation phenotype was highly variable across the species (**Fig. 1B** and **Fig. S2**), spanning all five score categories and covering more than eight orders of magnitude in actual transfer frequencies (**Fig. S1**; **Table S1**). Overall, strains are more often resistant than susceptible to conjugation. Only about one-third of strains (125/34) were consistently susceptible to both plasmids (score R388 >1 and score RP4 >1). Over 50% of the strains (n=189) were resistant to conjugation to at least one plasmid (R388 score <0.5 or RP4 score <0.5). 85 strains of them being resistant to both plasmids (score < 0.5), 82 strains were specifically immune to R388 and 22 were specifically immune to RP4.

Susceptibility showed a loose phylogenetic signal: clonal strains often shared the same score, yet closely related strains can differ markedly (**Fig. 1B**, lower-right inset). Immune and fully susceptible strains frequently occurred within the same sub-clades (**Fig. 1B**, lower-left inset), suggesting that factors in the accessory genome drive the phenotype. We thus hypothesized that susceptibility to conjugation could be affected by incompatibility with resident plasmids, known host defense systems, or prophages often carrying defense systems. To test this, we screened all genomes with DefenseFinder (*33*) and PADLOC (*34*) for known defense systems, and we identified prophages and plasmids using PHASTEST (*35*) and geNomad (*36*) (**Table S1**). Neither the presence/absence of specific defense systems nor the carriage of particular MGEs could be correlated with conjugation scores. An unbiased genome-wide association study likewise failed to reveal single genetic determinant with a robust effect. Our species-wide analysis demonstrates highly variable, dynamic, and plasmid-specific conjugation susceptibilities, indicative of fast-evolving barriers. The data suggest a multifactorial genetic basis, calling for forward-genetic screens to pinpoint the responsible determinants.

**Figure 1.**
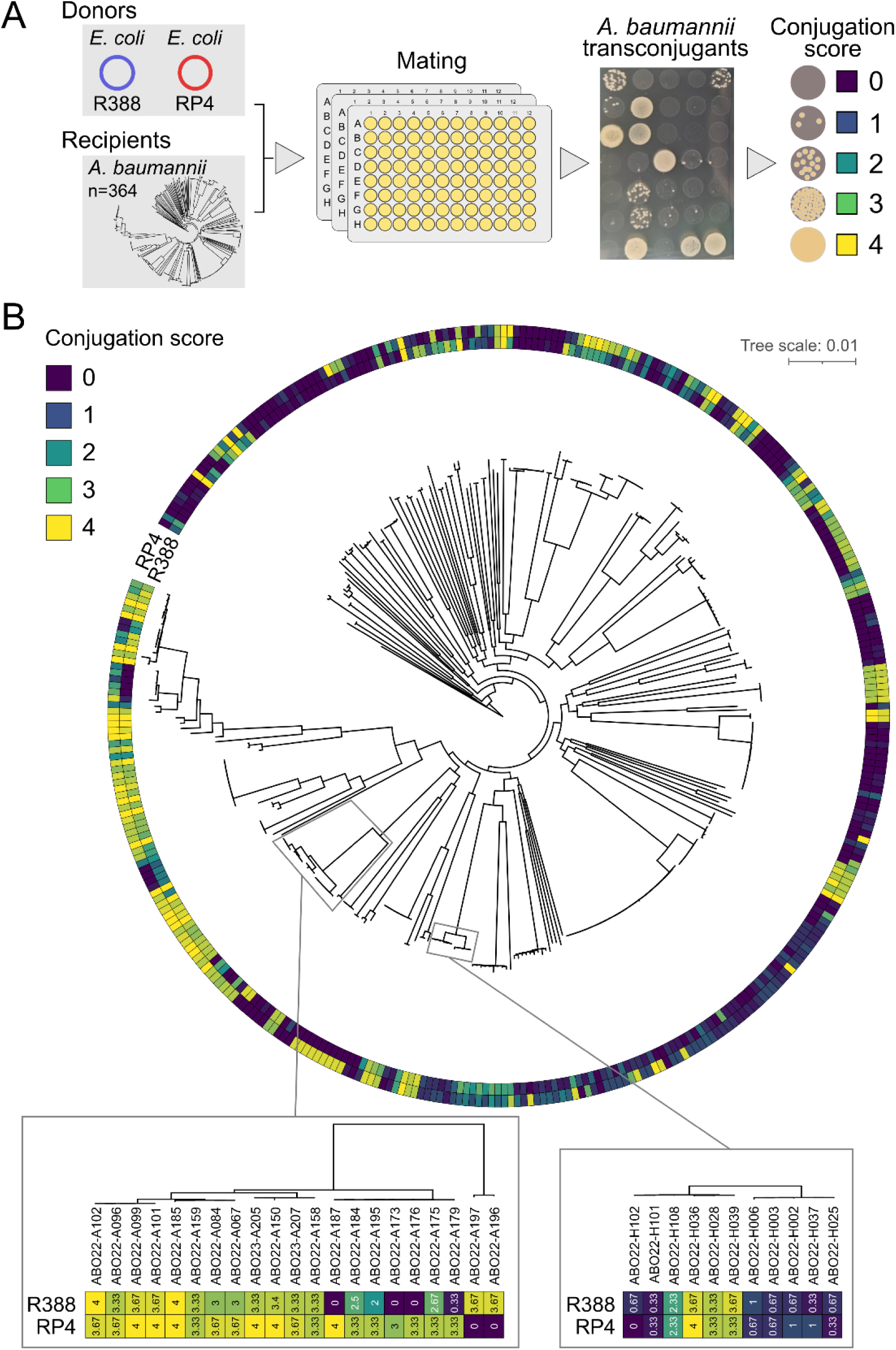
A high-throughput screening method reveals large variations in the susceptibility of diverse *A. baumannii* samples to receive plasmids R388 and RP4 by conjugation. (A) Overview of the species-wide conjugation score determination procedure. The *E. coli* donor strain BW25113 Δ*dapA*, carrying either R388 or RP4, was used in high-throughput mating assays with the 364 strains from the Ab-One collection. The number of transconjugants on selective plates was determined to associate a score for each strain, a reliable proxy of the actual conjugation frequency (**Fig. S1**). (B) Distribution of the conjugation scores for R388 (inner circle) and RP4 (outer circle) among the Ab-One collection presented as a rooted core-genome phylogeny (*32*). The scores for each strain, averaged from 5 independent experiments, are represented through a color heatmap, ranging from 0 (purple) to 4 (yellow). Insets show close-up on a subset of strains illustrating the strong variability within closely related strains. Tree scale is in substitutions per site.

### Genome-wide mutagenesis identifies strain-specific barriers to conjugation controlled by core genes

We hypothesized that inactivating genetic determinants conferring resistance to conjugation could restore plasmid acquisition, a phenotype that can be selected. To evaluate this, we sought to generate and screen Himar-1 transposon insertion libraries for mutants with increased susceptibility to conjugation. 166 strains with a score lower than 0.5 for at least one of the two plasmids were subjected to random mutagenesis. 107 strains failed to produce saturating mutagenesis (>10^5^ mutants), but we successfully obtained pools of >100,000 mutants for 17 non-clonal strains (**Fig. S2** and **Table S1**). Pools were subjected to conjugation by RP4 and/or R388 depending on the strains’ conjugation susceptibility. 12 mutant pools showed at least 10-fold higher conjugation frequencies than the non-mutagenized strains, indicating the presence of transposition insertions that suppressed resistance to conjugation. Surprisingly, sequencing of surviving colonies revealed recurring insertions into genes of the core genome (**Fig. 2A** and **Fig. S3**). From the 12 strains in which we identified transfer-restoring insertions, all but one (ABO21-A051) displayed hits in widely conserved genes. SbcD is the subunit of SbcCD nuclease that can cleave hairpin DNA (*37*), structures that are known to form on incoming ssDNA, notably to create hairpin promoters (*38*). Some strains show hits in *wzc* that could reduce the thickness of their capsule, possibly alleviating a physical barrier to conjugation (*27*). Other genes of metabolic pathways or encoding membrane-bound proteins, such as the highly conserved outer membrane transporter ZnuD (*39*), have so far no known role in regulating conjugation efficiencies.

Five different strains displayed multiple insertions upstream, but never within, the *hns* gene encoding the histone-like DNA-binding protein H-NS (**Fig. 2A**). We hypothesized that transcription coming from the transposon can increase *hns* expression, causing the restoration of conjugation. In three of the five strains in which the insertions were found (ABO22-A179, ABO22-A196 and ABO21-E077) we reconstructed a strong promoter upstream of *hns* and confirmed that this alleviated the strains’ resistance to conjugation (**Fig. 2B**). Deletion of *hns* in the overexpression mutant of ABO21-E077 (E077Δ*hns*) reverted conjugation frequencies to wild-type levels, proving that the effect is due to *hns* (**Fig. 2B** and **Fig. S4**). H-NS can act as a xenosilencing repressor, binding to AT-rich regions (*40*). It was previously found that recruitment of H-NS on the incoming plasmid depletes H-NS from the chromosome, causing a fitness cost associated with decreased conjugation efficiencies (*41*). We hypothesized that H-NS overexpression could overcome this effect. Yet, H-NS overexpression did not alter gene expression of an R388 derivative that could be introduced in strain ABO22-A179, suggesting that H-NS is not recruited on R388 (**Table S2**). Alternatively, and based on previous observations (*42*), we hypothesized that H-NS could restore conjugation by repressing an anti-plasmid activity conferred by a CRISPR-Cas. Yet, the expression of the type I-F CRISPR-Cas system of ABO21-A179 is not affected by H-NS overexpression (**Table S2**). Additional comparative RNAseq analyses of H-NS overexpression in strains ABO21-A051 and ABO22-A196, respectively suppressing barriers to R388 and RP4, show that these plasmid- and strain-specific effects are buried in a massive change in gene expression pattern (>600 genes in each strain, **Fig. S5** and **Table S2**). Further supporting a context-specific effect, the *hns*-overproducing promoter restored conjugation of R388, RP4 or both in 12 out of 39 additionally tested strains scattered throughout the species (**Fig. S2** and **Table S1**, see “hns_over”). Although H-NS has previously been reported to act as a barrier to horizontal gene transfer, this work reveals a more complex role in which H-NS can also promote conjugation through epistatic interactions with diverse determinants that limit plasmid acquisition.

**Figure 2.**
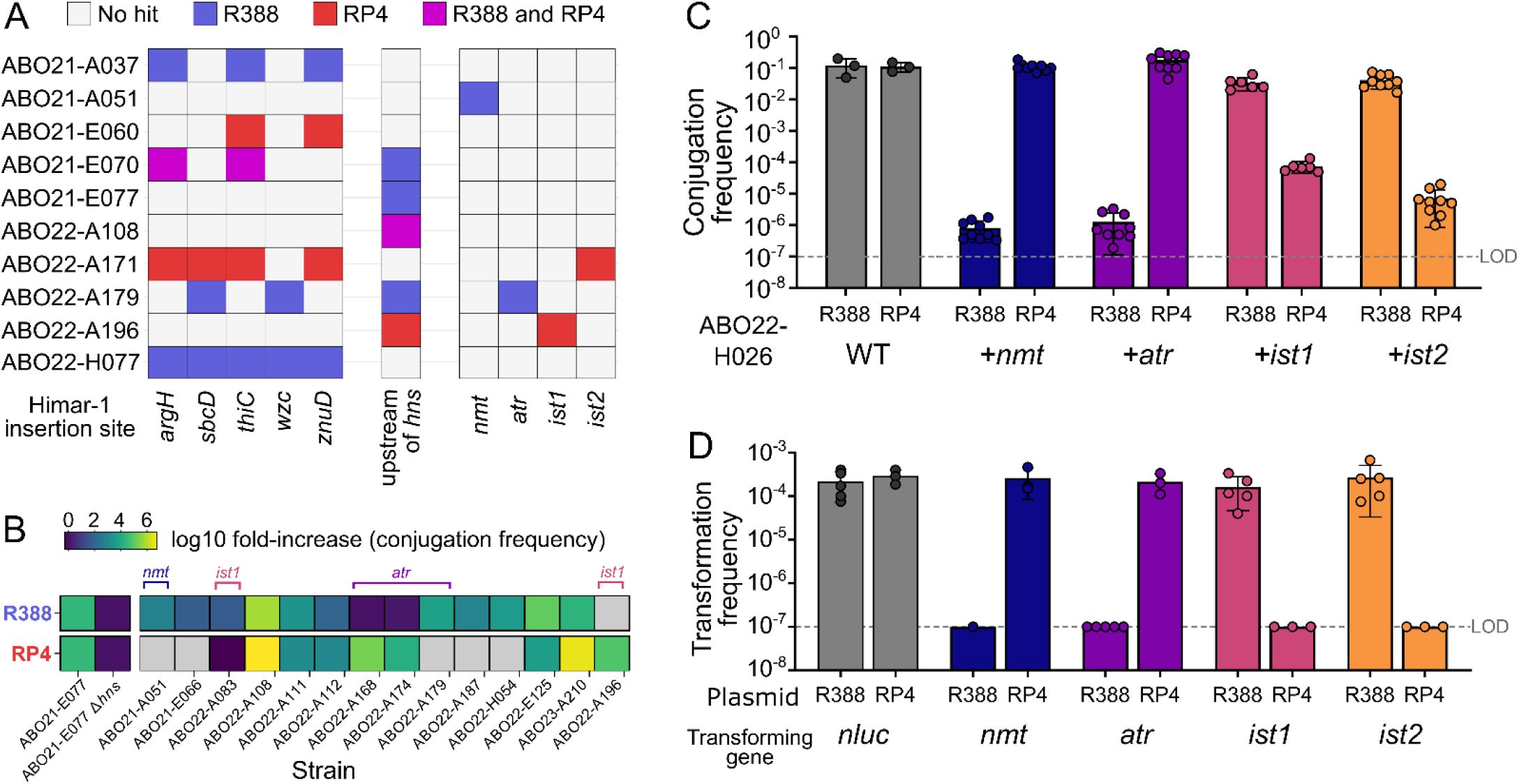
Random transposon mutagenesis identifies genetic elements conferring resistance to conjugation in a diverse subsample of *A. baumannii* strains. (A) Matrix of genes identified as inhibiting the transfer of R388 (blue), RP4 (red) or both (pink). Only the main hits are represented here, with the exhaustive list being depicted in **Fig. S3**. (B) Overexpressing *hns* in a diverse set of strains enhances plasmid transfer frequencies by multiple orders of magnitude. The value represented is the average fold-increase in transfer frequency (log10 scale, n=3). Grey cells represent situations of strains permissive to plasmids even without *hns* overexpression. Strains carrying one of the four genes conferring immunity to R388 and RP4 are represented with the name of the gene above the boxes. (C) Four genes encoding unknown function proteins confer plasmid-specific conjugation immunity when expressed in a heterologous but common genetic background (strain ABO22-H026, see **Fig. S7** for data in the ABO22-H008 strain). Bars represent the average of biological replicates (n=3-9), error bars represent the standard deviation. LOD: limit of detection. (D) Natural transformation of defense-encoding DNA in strains carrying R388 or RP4 displays the incompatibility between the defenses and their target plasmid. Individual points each represent a single replicate (n=3-5), with the bar end depicting average transformation frequency and SD represented with error bars. LOD: limit of detection.

### Four genes of patchy distribution confer specific immunity to R388 and RP4

In four strains, insertional inactivation of distinct genes, restoring conjugation of RP4 or R388 confirmed that barriers to conjugation are strain- and plasmid-specific (**Fig. 2A**). These four non-core genes were considered potential defenses against conjugative plasmids. Based on results presented in the next sections, we refer to them as members of the novel TAR (Targeted Anti-plasmid Response) class of defense systems. The genes acting against R388 were assigned to the Namtar and Attar sub-classes, respectively named *nmt* (ACT4ZX_13170) and *atr* (ACT4XV_16530). Those acting against RP4, were given the Ishtar sub-class and named *ist1* (ACT42K_09795) and *ist2* (ACT42V_03045). Namtar, Attar and Ishtar are ancient deities, respectively a Mesopotamian figure associated with death, a deity of the morning and evening star, and the goddess of love, war, and fertility.

Deletion/complementation of *nmt* and *ist1* in their original strain confirmed that the genes confer specific resistance to conjugation by R388 and RP4, respectively (**Fig. S6**). All four genes were then introduced in the *cysI* locus of strains ABO22-H026 and ABO22-H008, both permissive to R388 and RP4, and phylogenetically distant from those in which the genes were found (**Fig. S2**). In both strains, *nmt* and *atr* lowered conjugation frequencies of R388 by 5 orders of magnitude, without any effect on RP4 conjugation (**Fig. 2C** and **Fig. S7**). Conversely, *ist1* and *ist2* specifically decreased conjugation of RP4 by over 3 orders of magnitude (**Fig. 2C** and **Fig. S7**). Although *atr* and *ist1* genes were found in strains in which *hns* overexpression restored conjugation (**Fig. 2B**), their expression is not repressed by *hns* (**Table S2**) and *hns* overexpression in the strain ABO22-H026 carrying *ist*, *nmt* and *atr* genes did not suppress their protection against R388 and RP4 (**Fig. S8**). Thus, these genes function as independent monogenic immune systems against specific conjugative plasmids. While *ist1* is found in two clonal strains, *ist2* (n=13), *nmt* (n=5) and *atr* (n=24) are found in 4 to 6 independent lineages throughout the tree, suggesting horizontal gene transfer or repeated gene loss (**Fig. S2**). Further supporting that the genes are part of a dynamic repertoire, *ist1* is found in a variable region resembling an integron remnant, with other genes preceded by integron-type *attC* sites but lacking an integrase gene (**Fig. S9**). Similarly, *ist2* is found in between core genes, with various genes encoding unknown function proteins being shuffled at the same location (**Fig. S9**). On the other hand, *nmt* and *atr* are found in *loci* of high genetic diversity, with large regions not conserved between the strains. We also did not identify frequently co-localized genes that might contribute to their function. Hence, *ist, nmt* and *atr* encode standalone immune systems against conjugation.

### Ishtar, Namtar and Attar systems confer immunity against conjugative plasmids downstream of plasmid entry

We first sought to determine whether the *ist1, ist2*, *nmt* and *atr* genes prevented mating-pair formation or if they acted post-entry. Constitutive expression of *ist1, ist2*, *nmt* and *atr* genes respectively prevented transformation of R388 or RP4 delivered by electroporation (**Fig. S10**). Their retained activity and specificity toward the plasmid regardless of the mode of entry suggested that Ishtar, Namtar and Attar systems may be active post plasmid entry. To test this hypothesis, we took advantage that *A. baumannii* can import and recombine exogenous DNA in its chromosome by natural transformation. R388 and RP4 transconjugants of permissive strain ABO22-H008 were exposed to exogenous DNA carrying the *ist1, ist2*, *nmt* and *atr* genes along with a tetracycline resistance gene, all flanked by sequences allowing recombination in the chromosome. If active against the resident plasmid, bacteria transformed by the defense genes would not be obtained if selection pressure for the plasmid is applied. Indeed, while having no effect on the number of transformants obtained for the *nanoluc (nluc)* control gene, selection for R388 and RP4 resulted in a dramatic loss of *ist1, ist2*, *nmt* and *atr* transformants, respectively (**Fig. 2D**). Hence, Ishtar, Namtar and Attar systems can act on resident plasmids and can prevent formation of transconjugants by acting post plasmid entry. Yet, follow-up work shows that they differ in their mechanism of action.

### The widespread Ishtar immune system induces plasmid loss in RP4-carrying cells

The two Ishtar genes were identified separately on strains showing the same phenotype of being immune to RP4 conjugation. Although their primary sequence and different genetic environment suggested that they were distinct, the genes encode two proteins displaying a similar predicted structure (**Fig. 3A**). A C-terminal domain consisting of five or seven beta-sheets and three alpha-helices has no known function. Both proteins harbor a N-terminal HEPN (DUF4145 or DUF3644) domain, known to have a ribonuclease activity (*43*) with a demonstrated role in defense against phage (*44*, *45*) but which had to our knowledge no known anti-plasmid activity. Sequence homology searches across prokaryotic genomes revealed that Ishtar1 and Ishtar2 belong to a common family of defense proteins broadly distributed across sequenced prokaryotes (**Fig. 3B and Table S3**). Similar to their anti-phage counterparts, Ishtar-encoding genes are often found in the genomic proximity of other known defense genes. Like anti-phage systems they were also found recruited within integrons of *Vibrio* species and in P4 prophage satellites.

To further investigate their mode of action, the two *A. baumannii* Ishtar-encoding genes were placed under control of the anhydrotetracycline (aTc) inducible P_tet2_ promoter, and the RP4 plasmid was conjugated in those cells while keeping the Ishtar-encoding genes under repression. In the absence of RP4 selection, induction of the *ist* genes slowed growth (**Fig. 3C**) with an initial slight drop in viability based on CFU count but that remained constant over time (**Fig. 3D**). However, 5 minutes after induction, the percentage of bacteria forming RP4-carrying colonies decreased by over 8-fold, and by 40-fold at 30 minutes following *ist1* induction. Plasmid loss is even more dramatic following induction of *ist2* with over 99.5% of bacteria losing RP4 following 30 minutes of induction. Thus, the Ishtar system promotes rapid loss of RP4 with limited and transient effect on viability.

**Figure 3.**
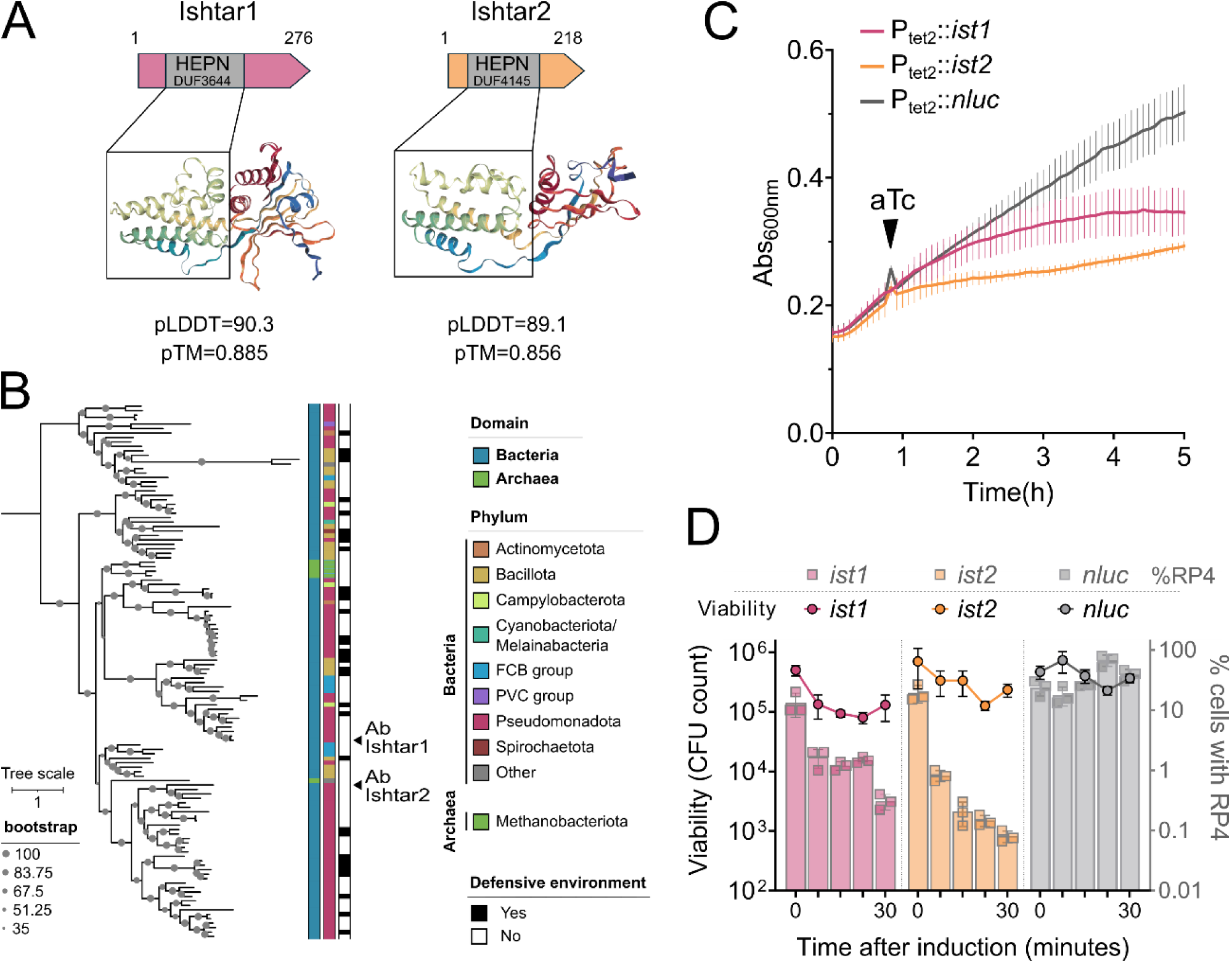
The Ishtar immune system induces plasmid loss in RP4-carrying cells. (A) Graphical representation of the protein domains (top) and structures (bottom) of the two members of the Ishtar class. HEPN domains are highlighted in grey and framed in grey rectangles on the 3D structures predicted with AlphaFold Server (AlphaFold 3) represented using NGLViewer (2.0.0-dev.37). (B) Distribution of homologs of the defenses in the tree of life, with the two original members of the Ishtar immune system highlighted with black triangles. Leaves were annotated as “defense-associated” when located within 10 genes upstream or downstream of a known defense system. (C) Induction of Ishtar defense systems in strains carrying RP4 slows bacterial growth as measured by absorbance (Abs_600nm_). Points (taken every 5 minutes) represent the average value for 3 replicates, SD represented with error bars. (D) Induction of *ist* genes in strains carrying RP4 has limited effect on cell viability (through total CFU counting) but show a rapid drop in plasmid-bearing population (% cells harboring RP4). For total CFU, points represent the average number of colonies for 3 replicates, SD represented with error bars. For % cells harboring RP4 (CFU with apramycin / CFU without selection for the plasmid), points each represent a single replicate (n=3), with the bar end depicting average value and SD represented with error bars.

### The transmembrane Namtar and Attar immune systems lead to the death of cells expressing the conjugative machinery

Namtar and Attar systems do not possess any characterized domain, but share the presence of a transmembrane region with five alpha helices (white striped bars) (**Fig. 4A**). Homology-based searches confirmed that Namtar and Attar belong to two distinct protein families found in hundreds of bacterial genomes belonging to diverse phyla (**Table S3**). Namtar homologs were detected in 0.47% of bacterial genomes with a broad taxonomic distribution, while Attar was found in 1.38% of bacterial genomes but was almost limited to Pseudomonadota (**Fig. 4B**). Notably, like Ishtar homologs, Attar homologs are also found in integrons of *Vibrio* species. Interestingly, several bacterial genomes encoded more than one Ishtar, Namtar and/or Attar homologs, suggesting that bacterial genomes can accumulate multiple anti-plasmid systems, like they do with anti-phage systems.

Contrasting with Ishtar, the expression of Namtar and Attar systems does not induce plasmid loss (**Fig. S11A**) but cause an immediate growth arrest (**Fig. 4C**) and a 100-fold (*nmt*) to 1,000-fold (*atr*) decrease in viable bacteria in the absence of plasmid selection (**Fig. 4D**). This is accompanied by a dramatic decrease in ATP levels, indeed supporting the conclusion that the presence of R388 drives Nmt and Atr to trigger an energetically depleted state and that cells are moribund (**Fig. S11B**).

During defense genes transformation experiments (**Fig. 2D**), we obtained a few *nmt* transformants in a strain bearing R388. We refer to these as “escapers” as they have evolved a way to escape the death-inducing Namtar defense. Sequencing of an escaper revealed a large deletion in R388, encompassing several *virB* genes encoding the conjugative apparatus. Indeed, deletion of the region encoding the MOB/MPF modules of R388 (*trwABC*+*virB2-11*, noted as Δtra) completely abolished the anti-R388 activity of Namtar, but also of Attar (**Fig. S12A**). Deletions of the two regions encoding the RP4 conjugation apparatus did not alter Ishtar activity (**Fig. S12A**). To further refine the requirements of Namtar and Ishtar, we sought additional rare transformants of *nmt* and *atr* in bacteria carrying R388. Sequencing revealed large and small deletions in the plasmids (**Fig. S12B**). All deletions in *nmt* transformants encompass *virB8*, with the smallest deletion of 8 bp (escaper3, esc3) creating a frameshift truncation at the C-terminus of VirB8. Deletions affecting *virB8* were not found in *atr* transformants carrying an R388, which instead showed a 118-bp deletion within *virB2* (escaper9, esc9) (**Fig. S12B**). We confirmed that the *virB8* and *virB2* deletion escaped Namtar and Attar, respectively (**Fig. 4E**). Supporting that *these* escapers were specifically selected by their associated defenses, a deletion of *virB8* could not escape Attar (**Fig. S12C**). Bacteria with R388 carrying the short deletion of *virB8* (esc3) and *virB2* (esc9) remained fully viable (**Fig. 4C and 4D**), indicating that growth arrest depends on Namtar and Attar respectively sensing VirB2 and VirB8. Hence, Namtar and Attar systems trigger a non-replicative and energetically depleted state in cell carrying R388. The growth arrest, dependent on specific VirB2 or VirB8 proteins of the conjugative system, is reminiscent of abortive infection response triggered by phage defense sensing phage proteins.

**Figure 4.**
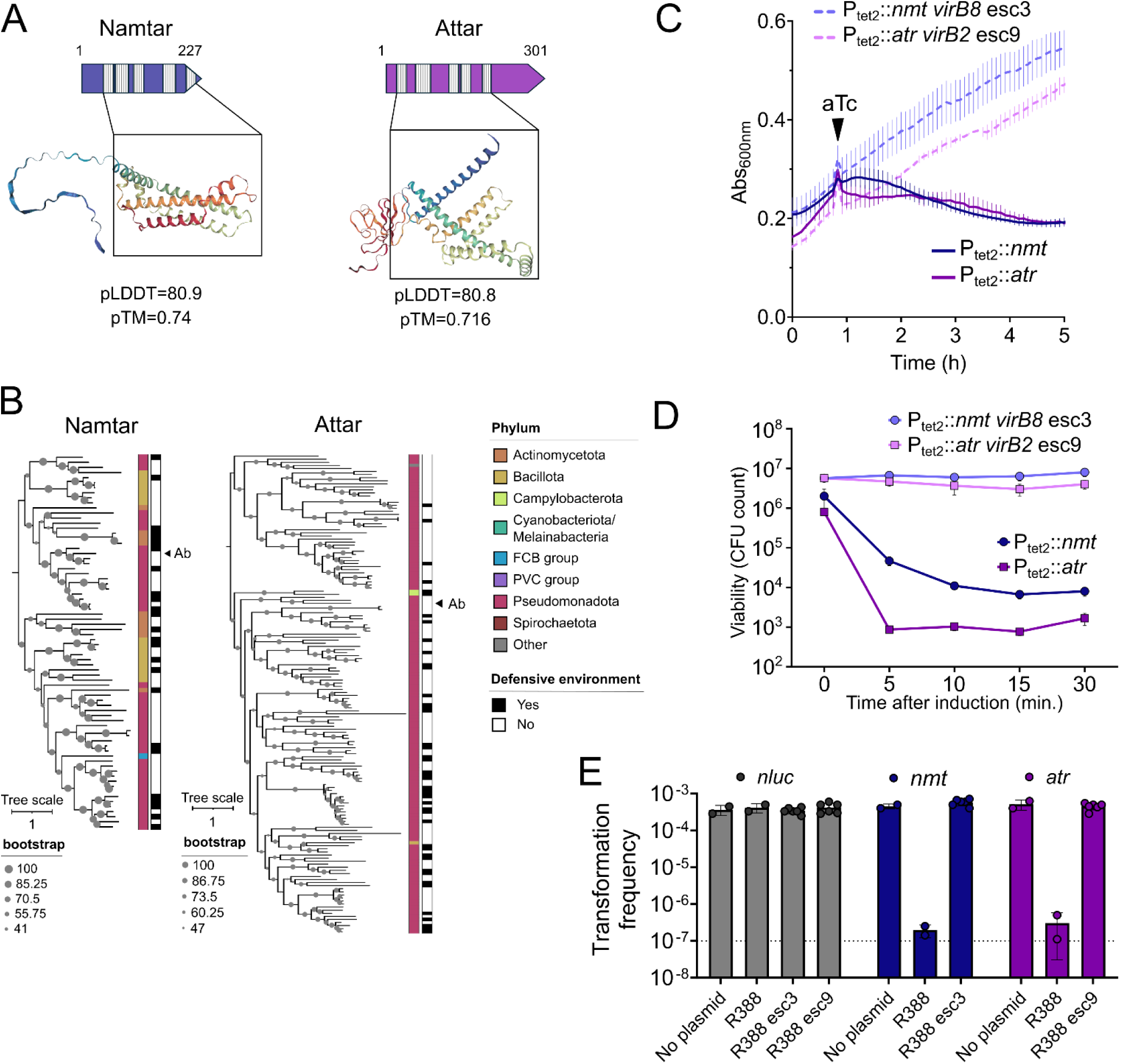
Namtar and Attar immune systems induce loss of viability in R388-carrying cells. (A) Graphical representation of the protein domains (top) and structures (bottom) Namtar and Attar. Vertical hashed lines represent predicted transmembrane regions, framed in grey rectangles on the 3D structures (predicted with AlphaFold Server (AlphaFold 3) represented using NGLViewer (2.0.0-dev.37). (B) Distribution of homologs of the defenses in the tree of life, with the original Namtar and Attar highlighted with black triangles. Leaves were annotated as “defense-associated” when located within 10 genes upstream or downstream of a known defense system. (C) Induction of Namtar defense systems in strains carrying R388 leads to a rapid Abs_600nm_ decrease, indicating cell death, in the presence of R388, but not in the presence of R388 escaper mutants. Points (taken every 5 minutes) represent the average value for 3 replicates, SD represented with error bars. (D) Induction of *nmt* and *atr* genes in strains carrying R388 causes cell growth arrest and loss of viability (through total CFU counting) in the presence of WT R388, but not VirB8 and VirB2 escaper mutants. For total CFU, points represent the average number of colonies for 3 replicates, SD represented with error bars. (E) Efficiency of natural transformation of the *nluc* (control), *nmt* and *atr* genes in strain carrying the escaper mutants of R388. Individual points each represent a single replicate (n=3-6), with the bar end depicting average transformation frequency and SD represented with error bars.

### Namtar confers population-level immunity by triggering a non-growing state in transconjugants

We identified a homolog of the *A. baumannii* Namtar in *Escherichia coli* (34% identity, WP_059327696.1, encoded by *nmt_Ec*). The two proteins share a similar structure (**Fig. S13A**), with only the N-terminal disordered region being different. To test if the function of Namtar is evolutionary conserved, we introduced the *nmt_Ec* gene or an *rfp* control into the chromosome of *E. coli* MG1655ΔRM (*46*) and measured the susceptibility of the resulting strains to conjugation of R388 and RP4. Transfers of R388 and RP4 to control RFP-expressing cells were highly efficient with nearly all recipients receiving the plasmid in 120 minutes (conjugation frequency of ∼1) (**Fig. S13B**). In contrast, *nmt_Ec* specifically decreased conjugation frequencies of R388 by over 4 orders of magnitude, recapitulating the observation made in *A. baumannii*, (**Fig. S13B**). In a time-course assay, Namtar caused a 1,000-fold reduction in the number of transconjugants relative to the RFP control within 30 minutes, a depletion reaching 10,000-fold after 120 minutes (**Fig. 5A**). Strikingly, while the population of RFP-expressing recipients increases, the population of Namtar-expressing recipients decreases, consistent with conjugation inducing a non-replicative moribund state in Namtar-expressing cells (**Fig. 5B**). To verify this, we sought to directly observe R388 conjugative transfer using timelapse microscopy and the ParB/*parS* reporter system (*47*). Briefly, the *parS* sequence from plasmid P1 was introduced into R388, which was transferred to recipient cells expressing the mChartreuse-tagged ParB_P1_ protein. Recipient cells present a diffuse green signal in the cytoplasm until the conjugative plasmid enters the cell and is converted into double-stranded DNA. This enables the recruitment of ParB_P1_ proteins to the *parS* sequence, resulting in the formation of bright green foci (**Fig. 5C**, green arrows). Tracking the fate of transconjugants, we observed that conjugative transfer of R388 blocked replication of transconjugants carrying *nmt_Ec* (**Fig. 5C**). Indeed, recipient cells expressing RFP (receiving or not R388) underwent 3 divisions on average during a 2-hour period (**Fig. 5D**). In contrast, while recipient cells expressing *nmt_Ec* but not receiving R388 behaved similarly, R388 transconjugants did not produce any offspring, or divided once if the process was nearly completed before conjugation occurred (**Fig. 5D** and **Supp. Movie 1**). Thus, Namtar homologs appear to function similarly in *E. coli* and *A. baumannii*. Although the Namtar system does not prevent entry of R388, it causes a drop in ATP levels and a non-reproductive state. While the latter would forbid horizontal transmission, the former blocks the plasmid vertical transmission, effectively preventing its propagation, conferring population-level protection.

**Figure 5.**
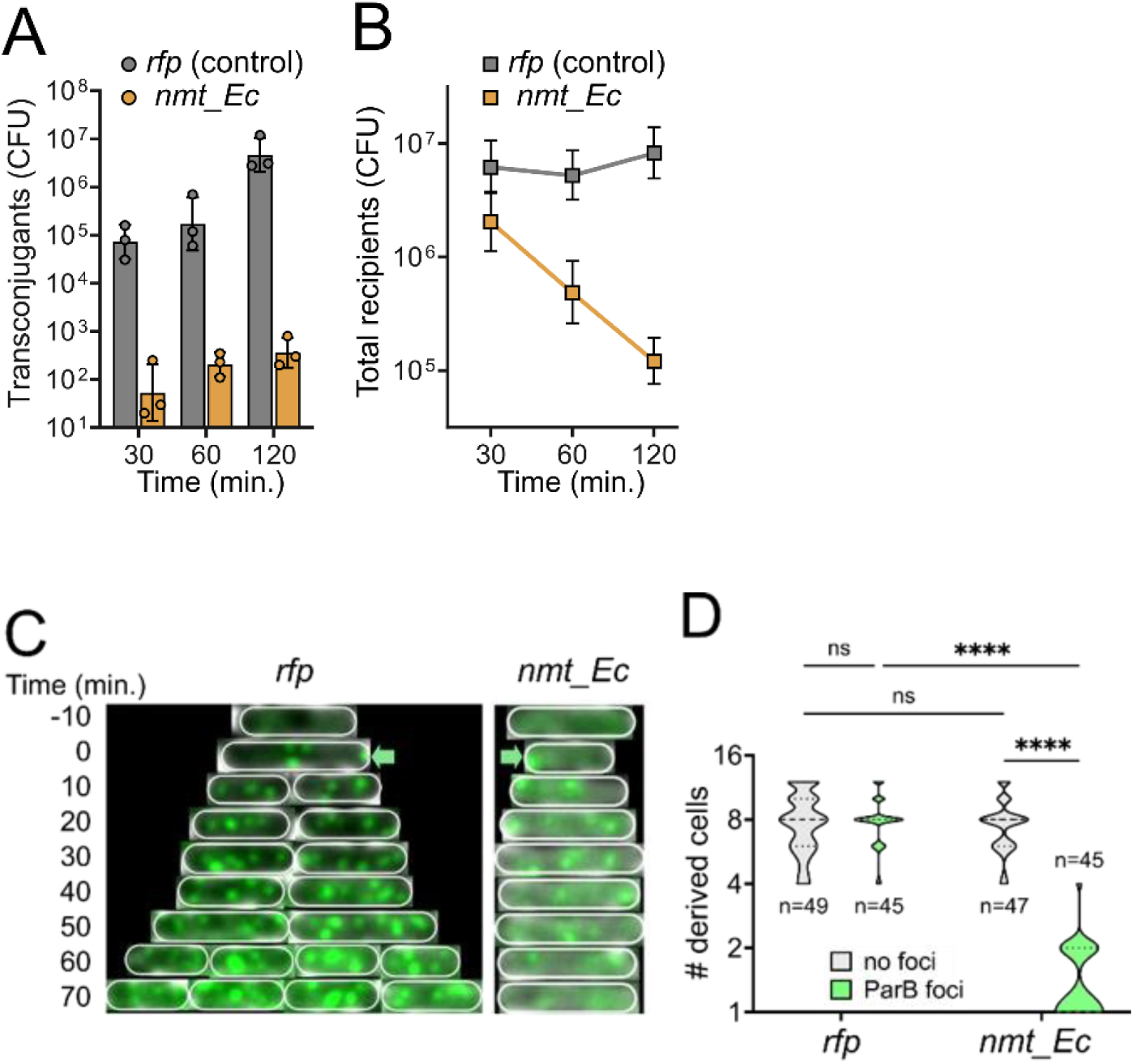
A functional Namtar homolog from *E. coli* induces a non-replicative state upon conjugative acquisition of R388. (A) Time course of conjugation efficiency, numbering transconjugants from mating BW25113 *dapA::aph* / R388 and recipient MG1655ΔRM derivatives expressing either a control *rfp* gene (coding RFP) or *nmt_Ec*. Conjugants CFU were determined by plating on LB + apramycin, with each point representing a biological replicate (n=3) and the bar end depicting average value. Standard deviation is represented with error bars. (B) Survival of the recipient population MG1655ΔRM expressing either a control *rfp* gene (coding RFP) or *nmt_Ec* during mating with BW25113 *dapA::aph* / R388. Recipients were determined by plating on LB. Points represent the average number of colonies for 3 biological replicates, SD represented with error bars. (C) and (D) Mating experiments in a microfluidic chamber, in which successful conjugation events were detected through the recruitment of ParB-mChartreuse to plasmid-borne *parS* sequences. (C) Kymograph of two representative recipient cells expressing RFP (left, “Control”) or *nmt_Ec* (right) during R388 acquisition. Plasmid entry is signaled by green arrows pointing towards the ParB-mChartreuse fluorescent foci and the outlines of cells (white). (D) Number of daughter cells derived from transconjugants expressing RFP (left, “Control”) or *nmt_Ec* (right) in the 120 minutes following the appearance of foci. The number of initial cells tracked is indicated. Statistical significance was tested using the unprotected Fisher’s LSD test from Graphpad Prism’s 2-way ANOVA (ns: p>0.05; ****: p<0.0001).

## Discussion

Our extensive assessment of conjugation in a species shows that barriers are not uniformly distributed and determined by the accessory genome. Although barriers could be alleviated by mutations affecting core genes - such as those causing increased H-NS expression - this may result from their interactions with accessory genes. H-NS is generally regarded as key maintainer of genome integrity, acting as a xenogeneic silencer of horizontally acquired genes (*40*), including of those encoding the conjugative apparatus (*41*, *48*, *49*) or directing transposons to AT-rich regions, away from essential genes (*50*). However, recent works are challenging this idea, with H-NS found to repress defense genes, possibly opening the gate to mobile genetic elements (*42*, *51*). The strain-specific alleviation of conjugation barriers may provide a means to uncover previously unrecognized barriers to conjugation. Interestingly, our functional approach did not reveal restriction-modification (R-M) as acting as conjugation barriers, although they are common in *A. baumannii* genomes (median=2; **Table S1**), suggesting that R-M systems represent a weak barrier to conjugative plasmids (*17*). However, this may also be attributable to the approach limitations and bias. Multiple strains could not be successfully mutagenized, possibly because of mechanisms that prevented transposon delivery by mobilization, such as R-M. This method could have also prevented us from identifying anti-conjugation systems that, for instance, detect conjugation attempts by sensing membrane perturbation or degrade incoming single-stranded DNA.

This approach nonetheless was powerful to identify four minimal, monogenic, plasmid-specific anti-conjugation genes. Members of the Ishtar (Incompatibility System via HEPN) immune system function by plasmid clearance. They appear specific to RP4, but we could not identify any escaper mutants pointing to a plasmid-specific protein. We currently favor the hypothesis that Ishtar acts by preventing the theta-type plasmid replication which depend on an RNA primer (*52*) and that could be degraded by the HEPN ribonuclease domain. Since R388 replicates using the same mechanism, this would imply that R388 carries a system to inactivate Ishtar, possibly in its leading region. This fits into an arms-race framework, with plasmids (or ICE) bearing anti-defenses to Ishtar, Namtar and Attar, possibly explaining that some strains carrying defense-encoding genes remain permissive to conjugation (**Fig. S2**). Members of the Namtar (Non-replicative/Abortive Mechanism) and Attar (Abortive Transfer) subclasses are two anti-R388 defenses which depend on VirB2 or VirB8, conferring specificity toward conjugative plasmids. Although they depend on distinct T4SS components, they both trigger a non-growing state with low ATP levels, possibly through membrane depolarization, that would ultimately lead to cell death. Their transmembrane segments suggest that they are membrane-bound sentinels capable of detecting the assembly of a conjugative system, signaling the presence of a conjugative plasmid that can spread through the population. Anti-conjugation defense triggering an abortive-like defense that sacrifices the host to prevent plasmid spread was unexpected. The identification of another VirB4-dependent anti-conjugation system in *E. coli* and also triggering cell death (*53*) supports that the abortive response is a common defense mechanism against conjugative plasmids. In contrast to virulent phages, conjugative plasmids do not pose a direct vital threat to the population. At best, they can incur a fitness cost but is often mitigated (*10*). However, conjugative plasmids can carry genes that may be ultimately harmful, such as gene-disrupting insertion sequences (*1*). The T4SS is also a common receptor to plasmid-dependent phage (*54*, *55*), and a population carrying a conjugative plasmid would be vulnerable to infection by such phages. Hence, an abortive infection-like response to acquisition of conjugative plasmids may have evolved through second-order selection, reducing the risk of future harmful events.

Phylogenetic and genetic context analyses showed that Namtar, Attar and Ishtar display features that are common in anti-phage defenses. Like some anti-phage defense, Ishtar carry an HEPN domain (*56*). Both Ishtar and Namtar can be found close to other defense genes, including in integrons where anti-phage defenses can also be compact and monogenic (*57*, *58*). Their widespread and patchy distribution suggests frequent gain and loss. Possibly because of structural and genomic context similarities, Namtar, Attar and Ishtar were classified as putative anti-phage proteins by a recent machine learning framework (*59*). However, the dependence of Namtar on the conjugative apparatus for activation argues against a role in phage defense. In addition, the *E. coli* Namtar did not confer protection against phages of the BASEL collection in our hands. Thus, current prediction models trained on anti-phage repertoires may also reveal anti-conjugation defenses.

In conclusion, our species-wide screen suggests that conjugative plasmids are subject to a potentially vast and diverse repertoire of host-encoded anti-conjugation defenses that have so far remained elusive. The identification of these mechanisms has important implications for understanding the spread and dissemination of antibiotic resistance by these mobile genetic elements.

## Materials and methods

### Bacterial strains, genetic modifications, and growth conditions

Bacterial strains, plasmids, and oligonucleotides used in this study are listed in Supplementary Tables S3, S4, and S5, respectively. *E. coli* BW25113 *dapA::aph*, auxotrophic for diaminopimelic acid, was obtained using to the lambda-red recombination method (*60*). Correct deletion of *dapA* was confirmed by PCR (DreamTaq Green Master Mix, Thermo Fisher Scientific) and Sanger sequencing (Microsynth, Vaulx-en-Velin, France). Genetic modifications of *A. baumannii* were performed by overlap extension PCR to synthesize a chimeric DNA fragment flanked by 2-kb homology arms targeting the insertion site, as previously described (*61*). Briefly, the fragment (2 ng/µL) was added to 200 µL of transformation medium (TM, tryptone medium supplemented with 0.6 mM CaCl_2_ and 0.6 mM MgCl_2_) (*62*) inoculated with 2 µL of an overnight culture of the strain to be modified. The suspension was incubated overnight at 37°C, after which half the volume was plated on selective medium supplemented with the appropriate antibiotic. Strains were grown in lysogeny broth (LB; Lennox formulation) or tryptone medium (5 g/L Bacto Tryptone, Gibco). Unless otherwise stated, all experiments were performed at 37°C in 2 mL of medium in 13-mL culture tubes. Where required, the following supplements were added at the indicated concentrations: ampicillin (Roth, K029.4), 100 µg/mL; anhydrotetracycline (aTc; Takara, 631310), 200 ng/mL; apramycin (Thermo Fisher Scientific, J66616.03), 60 µg/mL; chloramphenicol (Sigma-Aldrich, C3175), 25 µg/mL; hygromycin (Roth, 1287.1), 200 µg/mL; kanamycin (Roth, T832), 50 µg/mL; tetracycline (Fisher, 10460264), 5 µg/mL; diaminopimelic acid (DAP; Sigma-Aldrich, 33240), 0.3 mM.

### Modifications of conjugative plasmids for use in *A. baumannii*

The BW25113 dapA::*aph* donor strain was transformed by R388 and RP4 using electroporation. To detect their transfer to *A. baumannii* strains, an apramycin resistance marker (*aac4*, amplified from pASG1 (*63*) was inserted in place of the *intI1* integron for R388 and within the ampicillin resistance gene *bla* for RP4, using the lambda-red method (*60*). Along with *aac4*, the coding sequence of NanoLuc® luciferase with a modified promoter and added *ssrA*/TEV degradation sites was also inserted. The resulting plasmids are named RP4-star and R388-star but referred to R388 and RP4 in the manuscript. Plasmids were sequenced using Oxford Nanopore Technology. R388* and RP4* plasmids are available from Addgene under catalog numbers 258681 and 258682, respectively. Variants of R388 (*Δtra*) and RP4 (*Δtra1*, *Δtra2*, *tetAR:*:*aac*, Δ*tetAR*, Δ*aph*, Δ*aph*-*tetAR*::*aac*) were constructed directly in *A. baumannii* strain ABO22-H026 using the method described in previous section.

### Determination of conjugation rates

Two different methods were used depending on the number of conditions to assay and the required accuracy of conjugation efficiency. Method 1, a high-throughput semi-quantitative method used to screen the *A. baumannii* collection, relies on the use of 96-well plates. Donors BW25113 *dapA*::*aph* carrying RP4 or R388 were grown overnight at 37°C in flasks; recipients were grown overnight at 37°C in 200 µL LB in 96-well plates. The next day, donors were pelleted and resuspended in the same volume in LB with 0.6 mM DAP and mixed in a 1:1 ratio with cultures of the recipients. 96-well plates were prepared by dispensing 150 µL of molten agar LB-DAP (0.3 mM) in each well to form flat, bubble-free surface. Once the plates were dried, 20 µL of each suspension were mixed in a separate 96-well plate, then 20 µL were dropped on top of each well. Plates were let to dry for 1 hour, then incubated overnight at 37°C, upside down. Cells were then recovered by dispensing 100 µL of LB on top of each well and repeated pipetting. The resulting suspension was transferred to a new 96-well plate. 10-µL drops spotted onto LB with apramycin were used to determine conjugation scores as defined in the result section (**Fig. 1A**). This method was also used to measure conjugation frequencies by performing serial dilution of the suspension and numbering total recipients and transconjugants. The frequency of conjugation is reported as the number of transconjugants divided by the total number of recipients. Method 2, the conventional quantitative method, was used to obtain conjugation rates. Donors and recipients from overnight cultures were pelleted and resuspended in LB-DAP in 1/10^th^ of the original volume. Cells were mixed in a 1:1 ratio (50 µL of each suspension) before spotting 80 µL drops on LB-DAP agar plates. Once dried, the plates were incubated at 37°C for 4 hours. Then, bacteria were scraped from their plates using a sterile loop, resuspended in 1 mL LB, pelleted, and resuspended in 200 µL of LB. This suspension was then serially diluted (down to 10^-7^), and 10 µL drops were spotted on the appropriate selective media before overnight incubation at 37°C. The frequency of conjugation is reported as the number of transconjugants divided by the total number of recipients.

### Random mutagenesis by transposition and selection of conjugation-permissive mutant

Mutants were generated using the suicide vector pSC189 carrying a Himar-1 transposon conferring kanamycin resistance (*64*). The *E. coli* MFDpir donor strain (*65*) carrying pSC189 is mated with the strain for 2 hours at 37°C, on LB-DAP agar plates, in a procedure identical to the one followed for determining conjugation rates. Then, cells are scraped, resuspended in sterile LB, and spread onto LB plates with kanamycin to select transposition mutants. Mutant libraries of 100,000 mutants were constructed as 10 sub-pools, each composed of 10,000 mutants. Sub-pools were individually tested for conjugation using the previously described high-throughput protocol and the number of conjugants obtained was compared to the wild-type strain. Transconjugants obtained were then either isolated individually or pooled altogether to identify the insertion sites.

### Identification of transposon insertion sites

Mutants bearing a transposon insertion that restored the ability of the cell to receive a plasmid were collected, and their genomic DNA was extracted using QIAGEN’s DNeasy Blood & Tissue Kit (QIAGEN, 69504). Then, the inverse PCR (iPCR) technique was used to identify transposon insertion sites. Briefly, genomic DNA was digested with AseI (New England Biolabs, R0526S) or DraI (New England Biolabs, R0129S), then fragments were self-ligated with T4 DNA ligase (New England Biolabs, M0202S). Finally, iPCR was used to amplify the insertion site using primers binding within the transposon sequence, going in opposite directions towards the transposon’s boundaries. For individual mutants, amplicons sequences were mapped onto the reference genome to identify the transposon insertion site using the SnapGene software. For pools of mutants, amplicons were sequenced using Oxford Nanopore Technology. Mapping of the reads onto the reference genome was done using InMut-finder (*66*). Genes were noted as hit when at least 20 reads mapped to the same location. When multiple insertion sites were observed for a single gene, at separate locations, they were grouped.

### Heterologous expression of candidate defense genes at the *cysI* locus

Heterologous chromosomal expression platforms were constructed using transformation of overlap extension PCR fragments as previously described. Constitutively expressed (*tetA*-gene of interest) and inducible (*hph*-*tetR-*Ptet2-gene of interest) versions were constructed for each candidate (using the *nluc* gene as control) in strain ABO22-H026. They were then re-amplified and transformed into other strains when needed. After sequencing each mutant (Sanger sequencing, Microsynth, Vaulx-en-Velin, France) to verify that the sequences were correct, various assays were performed according to the expected observations. Accession numbers for the 4 members of the TAR family are the following: XZU70292.1 (*nmt*), MGQ1295451.1 (*atr*), XZW63086.1 (*ist1*) and MGQ1310977.1 (*ist2*).

### Natural transformation of defense genes in plasmid-bearing strains

DNA fragments containing the modified *cysI* locus carrying the defense genes, or control, under constitutive expression driven by tetracycline resistance gene *tetA* were amplified along with 2-kb flanking regions on each side. These fragments were then naturally transformed, as previously described, into strains bearing either R388, RP4, their respective variants, or no plasmid. After overnight incubation in TM, the suspension was serially diluted, and 10-µL drops were spotted on selective media to quantify the total number of bacteria and the number of transformants. Transformation frequency was defined as the number of transformants divided by the total number of bacteria. Each experiment was performed in triplicate.

### Determination of the impact of defense induction in plasmid-bearing strains

ABO22-H026 variants harboring the genes of interest under the control of the aTc-inducible promoter Ptet2 (*67*) were grown overnight at 37°C with shaking, before being diluted 1/50^th^ in LB containing apramycin (to ensure plasmid maintenance among the population) and incubated at 37°C with shaking for around 3 hours. Upon reaching an optical density (600nm) of 0.5, cells were pelleted and resuspended in LB and 150 µL of each suspension was transferred to a 96-well plate. Supplements (apramycin) were then added to the appropriate wells, and the plate was incubated for 40 minutes at 37°C in a plate reader (Infinite M200 PRO, Tecan Group Ltd., Switzerland) while monitoring Abs_600nm_. After this growth period (start of the exponential phase, visible increase in bacterial density on the curves), plates were taken out and aTc was added to the appropriate wells, before putting the plates back in the plate reader to monitor Abs_600nm_ for the rest of the experiment. For CFU counting and ATP measurements, a second identical plate was set up in the exact same conditions and taken out of the plate reader at various timepoints, to ensure that the absorbance measurement would not be affected by repeatedly opening the reader and the plate’s lid. For each timepoint (0, 5, 10, 15, 30 minutes post-induction), a sample was taken out of the second plate, and to quantify ATP levels and determine viability by CFU counts. Samples were immediately used for ATP quantification by mixing 20 µl of bacterial suspension with an equal volume of BacTiter-Glo™ reagent (Promega Corporation, Madison, WI, USA), and luminescence was immediately measured using the GloMax® Navigator luminometer (Promega Corporation, Madison, WI, USA). Luminescence Units (LU) were normalized to the measured cell density via Abs_600nm_, yielding Relative Luminescence Units (RLU). To determine viable bacteria, samples were serially diluted after being taken out of the plate reader, and 10-µL drops were spotted onto media to select all live bacteria (on LB), and live bacteria still harboring the plasmid of interest (on LB with apramycin). Plates were incubated overnight at 37°C and CFU were counted to quantify each population.

### Construction of *E. coli* strains carrying *nmt_Ec*

The genes *nmt_Ec* (encoding WP_059327696.1) or *rfp* (encoding RFP) were introduced in MG1655ΔRM in the attTn7 locus of the strain, using a 2-plasmid strategy (*68*). Briefly, the gene of interest is cloned on a suicide vector (R6K ori, kanamycin resistance) that harbours a cargo (spectinomycin resistance gene plus gene of interest) contained between two Tn7 recombination arms, in the E. coli strain DH5α λpir. Then, a helper plasmid (thermosensitive pSC101 ori, ampicillin resistance) carrying the Tn7 transposase genes (under the control of an arabinose-inducible promoter) is introduced in the strain. After arabinose induction, the suicide vector is electroporated into competent cells, which are left to recover for an hour, before plating on LB plates with spectinomycin. Helper and suicide plasmid loss was verified by serial streaking colonies on LB with ampicillin and LB with kanamycin, and only double sensitive clones were kept. Correct insertion was verified by PCR amplification of the attTn7 locus and Sanger sequencing (Microsynth).

### Live-cell visualization of conjugation and fate of the recipient cells

The *parS* sequence from plasmid P1 paired to a kanamycin resistance gene was introduced in place of the *intI1* integrase of R388 using natural transformation in *A. baumannii*. The resulting plasmid, named R388_*parS*_P1, was then introduced in the donor strain BW25113 by electroporation. MG1655ΔRM carrying *nmt_Ec* or *rfp* were electroporated with pNTM01-*parB*P1-mChartreuse, to track plasmid transfer in recipients as previously described (*69*). For live-cell imaging, overnight cultures were performed in M9 medium supplemented with glucose (0.2%) and casamino acid (0.4%) (M9-CASA) supplemented with kanamycin (for donor cultures) and ampicillin (for recipient cultures) at 37°C. The following morning, they were diluted to an OD_600nm_ of 0.05 in M9-CASA medium and were grown to an OD_600nm_ of 0.8 at 37°C. After washing off antibiotics, cells were resuspended in an equal volume of M9-CASA medium before being mixed in a 1:1 donor:recipient ratio. The different mixes (recipients expressing different genes of interest) were then loaded into a B04A microfluidic chamber (ONIX, CellASIC®) (*70*). Nutriment supply was maintained at 6.9 kPa and the temperature maintained at 37°C for the duration of time-lapse imaging. Cells were imaged every 10 min for 5 hours.

Conventional wide-field fluorescence microscopy imaging was conducted on an Eclipse Ti2-E microscope (Nikon), equipped with X100/1.45 oil Plan Apo Lambda phase objective, ORCA-Fusion digital CMOS camera (Hamamatsu), and using NIS software for image acquisition. Acquisitions were performed using 50% power of a Fluo LED Spectra X light source at 488 nm wavelength. Exposure settings were 500 ms for mChartreuse, and 50 ms for phase contrast. Images were then processed using Fiji software (*71*) with the MicrobeJ plugin (*72*), to extract quantitative data about the fate of conjugants originating from different recipient backgrounds. To do so, conjugation events were highlighted by the appearance of a green focus in recipient cells. The final number of cells derived from these conjugants was manually monitored for 120 minutes. As a control, an approximately equal number of recipient cells not having received the plasmid (no foci) were randomly chosen and treated in the same way.

### Gene expression analysis and expression profiling by RNAseq

Bacteria were prepared similarly to the way they would be used as recipients in conjugation assays. Briefly, after an overnight culture, cells were washed in an equal volume of LB, then 100 µL drops were spotted onto LB-DAP medium. The dried plates were then incubated for 4 hours at 37°C. Cells were then scraped from the plate and resuspended in LB before removing the supernatant. The resulting pellets were then stabilized and lysed as previously described (*73*). These samples were subsequently treated with DNase I and purified. Ribosomal-depleted RNA was used to construct strand-specific libraries, sequenced using Illumina NovaSeq X Plus Sequencing System (Novogene GmbH, Germany). Raw read processing and differential expression analyses were carried out using the Curare pipeline (*74*) for differential gene expression analysis. Raw reads are available at NCBI under Bioproject Accession number PRJNA1456853.

### Phylogenetic analyses and *in silico* characterization of candidates

Phylogenetic analyses were conducted on protein sequences from 41,540 complete bacterial genomes and 600 complete archaeal genomes downloaded from Refseq in August 2024. Known defense systems were predicted using DefenseFinder (v2.0.0) (*33*). Bacterial and archaeal sequences were separately filtered for redundancy using the clusthash function from MMseqs2 (*75*) (identity>99% and identical length). Representative non-redundant proteins were combined into a database of 49,315,157 sequences, which were further clustered into 3,394,584 clusters with MMseqs2 (parameter c -0.8). Each anti-plasmid protein was searched against these clusters using blastp (*76*), and non-redundant proteins from the highest-ranking cluster were aligned with Clustal-omega (*77*) to generate a seed HMM profile with hmmbuild (*78*). Seed HMM profiles were searched into non-redundant sequences with hmmsearch (*78*). Hits passing selection threshold (e-value <10^-5^ and query coverage >70%) were selected and domain sequences were collected based on envelope residues. Domain sequences were clustered with the easy-cluster function of MMseqs2 (--min-seq-id 0.9 -c 0.8), and representative sequences were aligned with MAFFT (*79*). A new HMM profile was generated from the MAFFT-generated MSA. This procedure was repeated twice. The final MSA was trimmed with clipkit (*80*) (default parameters) and a prokaryotic phylogenetic tree was computed with IQ-tree2 (*81*), with models VT+F+G4, LG+F+I+G4 and LG+F+G4 selected by ModelFinder for *nmt*, *ist1/ist2* and *atr* respectively. Node support was computed using 1000 iterations of the ultrafast bootstrap function in IQ-tree2 (option -bb 1000). All trees were visualized with iTOL. Leaves were annotated as “defense-associated” when located within ten genes upstream or downstream of a known defense system.

## Supporting information

Table S3. Immune systems homologs

Table S6. Oligonucleotides

Table S5. Plasmids

Table S4. Strains

Table S1. Conjugation score and metadata

Table S2. RNAseq analyses

## Acknowledgements

Plasmids R388 and RP4 were gifts from Sarah Bigot (MMSB, Lyon, France). Plasmid pKD46 and its variant harboring gentamicin resistance pKD46-Gm were gifts from Benoît Doublet (INRAE, Nouzilly, France). We thank Adrien Ducret (CIRI, Lyon, France) his helpful advice on microscopy image analysis. We thank Manon Bouvier for providing the *parS* sequence to modify R388. We acknowledge the occasional use of large language models to improve text clarity and concision.

## Author contributions (CREdiT)

LP: Investigation, Data curation, Formal analysis, Visualization, Writing – original draft, Writing – review & editing

JB, SG, BC: Investigation

JC, CL: Resources

FR: Investigation, Validation, Supervision, Funding acquisition, Writing – review & editing

XC: Conceptualization, Data curation, Investigation, Validation, Supervision, Funding acquisition, Writing – review & editing

