## Supplementary figures and images for "Widespread immune systems protect bacteria against conjugative plasmids"

### Supplementary movie S1

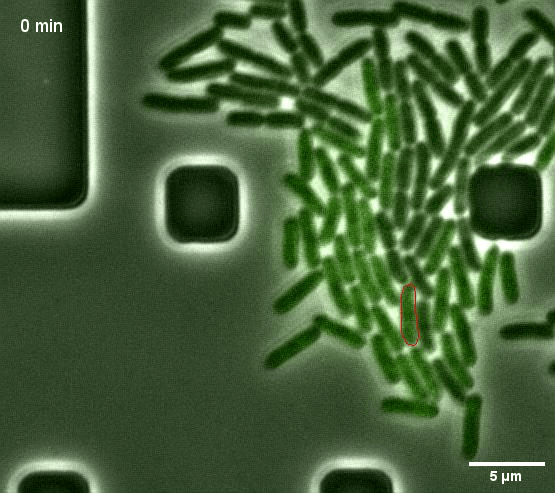
