## Supplementary Figures 1 to 13 for "Widespread immune systems protect bacteria against conjugative plasmids"

**This document contains :**

**Supplementary figures:**

**Figure S1.** Correlation between the observed conjugation score and measured conjugation frequency.

**Figure S2.** Phylogeny of the Ab-One collection with additional metadata describing the presence of defense homologs, results of the random mutagenesis screen and *hns* overexpression.

**Figure S3.** Exhaustive list of hit candidate genes identified as regulating plasmid conjugation to recipients.

**Figure S4.** Impact of *hns* overexpression on conjugation frequency in multiple strains.

**Figure S5.** Number of differentially expressed genes after analysis of the transcriptomic profile of *hns* overexpression mutants.

**Figure S6.** Confirmation of the role of *nmt* and *ist1* as barriers to conjugation, in their original strains through deletion and complementation.

**Figure S7.** Confirmation of the role of the four defenses on conjugation, through heterologous expression in the model strain ABO22-H008.

**Figure S8.** Effect of H-NS overexpression on conjugation in strain ABO22-H026 expressing the four conjugation defenses.

**Figure S9.** Genetic environment of the four anti-conjugation defenses.

**Figure S10.** The four defenses are active against conjugative plasmids delivered by electroporation.

**Figure S11.** Namtar and Attar defenses cause *virB8*- and *virB2*-dependent cell growth arrest and ATP depletion.

**Figure S12.** Deletions in *virB8* and *virB2* allow R388 to escape *nmt* and *atr*, respectively.

**Figure S13.** Structural and functional similarities between Namtar homologs in *A. baumannii* and *E. coli*.

**Supplementary tables:**

**Table S1.** Metadata for each strain of the *A. baumannii* Ab-One collection

**Table S2.** RNAseq analyses of mutants with *hns* overexpression

**Table S3.** HMM search results of Ishtar, Mantar and Attar immune systems

**Table S4.** Strains used in this study

**Table S5.** Plasmids used in this study

**Table S6.** Oligonucleotides used in this study

### Additional file not in this document:

**Supplementary Movie S1.** Time-lapse movie of cell growth arrest following the entry of R388 plasmid in recipient cells expression *nmt_Ec*.

Related to the figures 4C-D. Time lapse microscopy of a recipient cells expressing *nmt_Ec* who stops growth after R388 plasmid entry revealed by ParB focus formation. Cells were grown in M9-CASA at 37°C, and images were acquired every 10 minutes. Scale bar, 5 µm; time is indicated in minutes. Donor cells were in black and recipient cells were in green. Red arrows point to the fluorescent foci signaling plasmid entry in the recipient cells, red circles were drawn around cells of interest

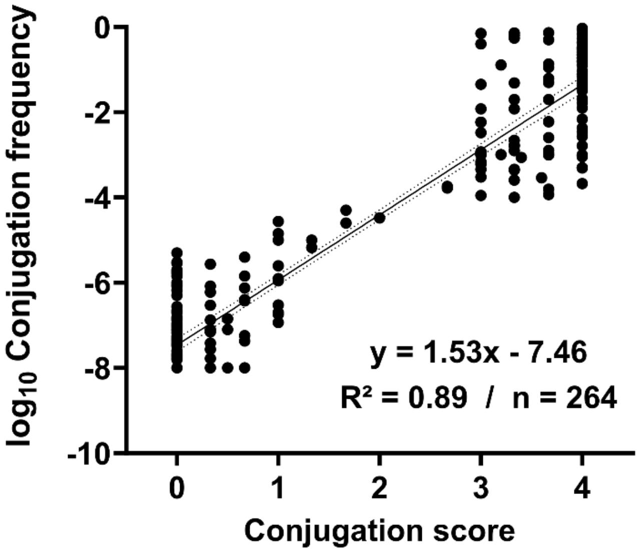


**Supplementary Figure S1.** Correlation between the observed conjugation score and measured conjugation frequency. For each of the 264 tested strains, the high-throughput mating assay was used, and the two metrics (log10-frequency vs. score) were compared using a simple linear regression ($R^2$=0.8926, p <0.0001).

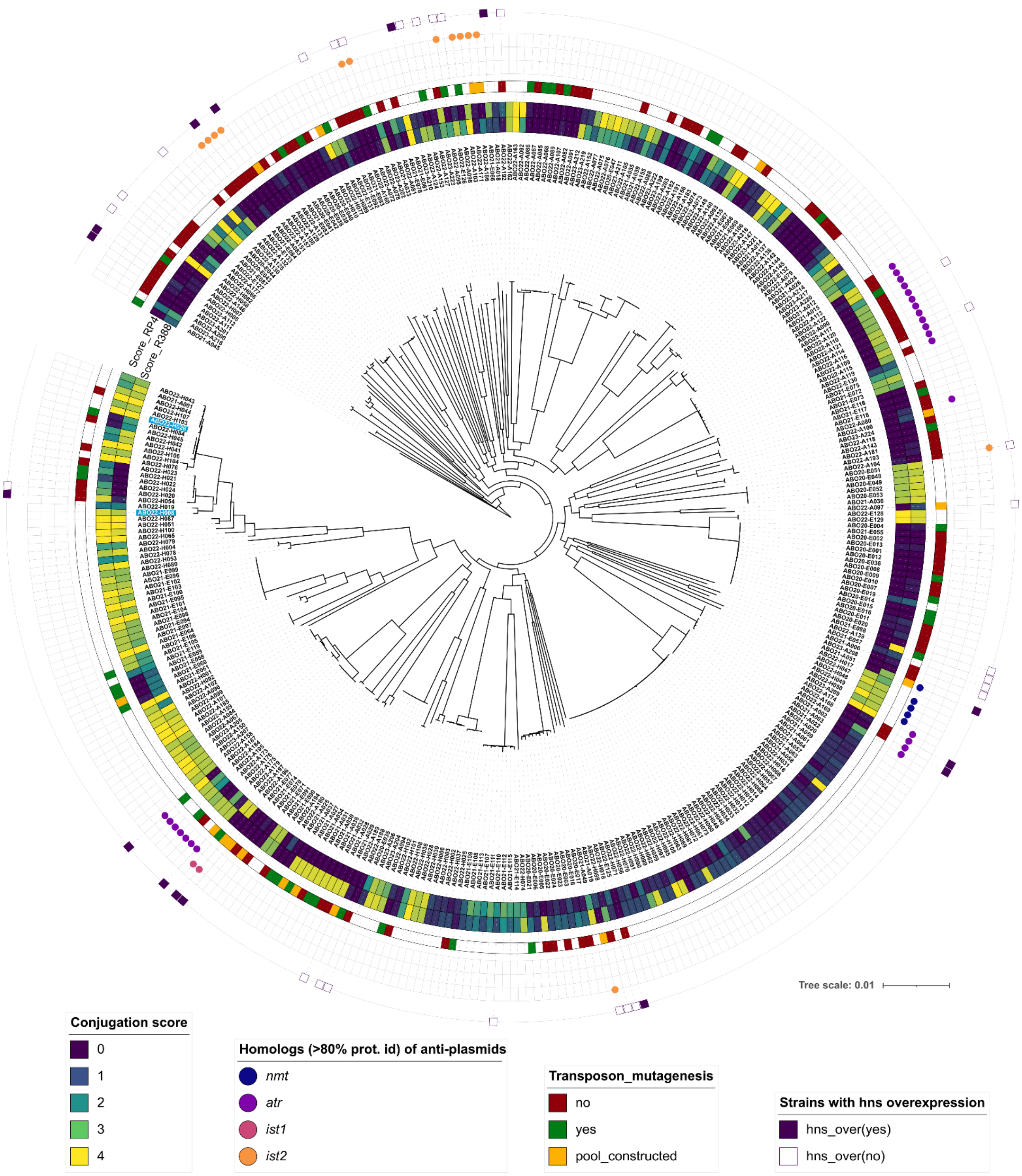


**Supplementary Figure S2.** Phylogeny of the Ab-One collection with additional metadata describing the presence of defense homologs, results of the random mutagenesis screen and hns overexpression. Distribution among the Ab-One collection of: 1) the defense proteins and their sequence homologs (colored dots), 2) strains submitted to random mutagenesis (red = not enough mutants generated; green = enough mutants, process not led to the end; yellow = pool constructed and tested for conjugation), 3) strains submitted to *hns* overexpression, with the results (no square = not tested; white square = no effect; purple square = plasmid transfer frequency increased by at least 10-fold). The strains used for heterologous defense expression (ABO22-H008 and ABO22-H026) are highlighted in blue. Phylogeny was built with IQTree2 using GTR+F+I+G4 (identified by ModelFinder), 1000 ultrafast bootstraps (--ufboot 1000) and UFBoot's tree optimization (--bnni). Visual representations were built using iTOL. Tree scale is in substitutions per site.

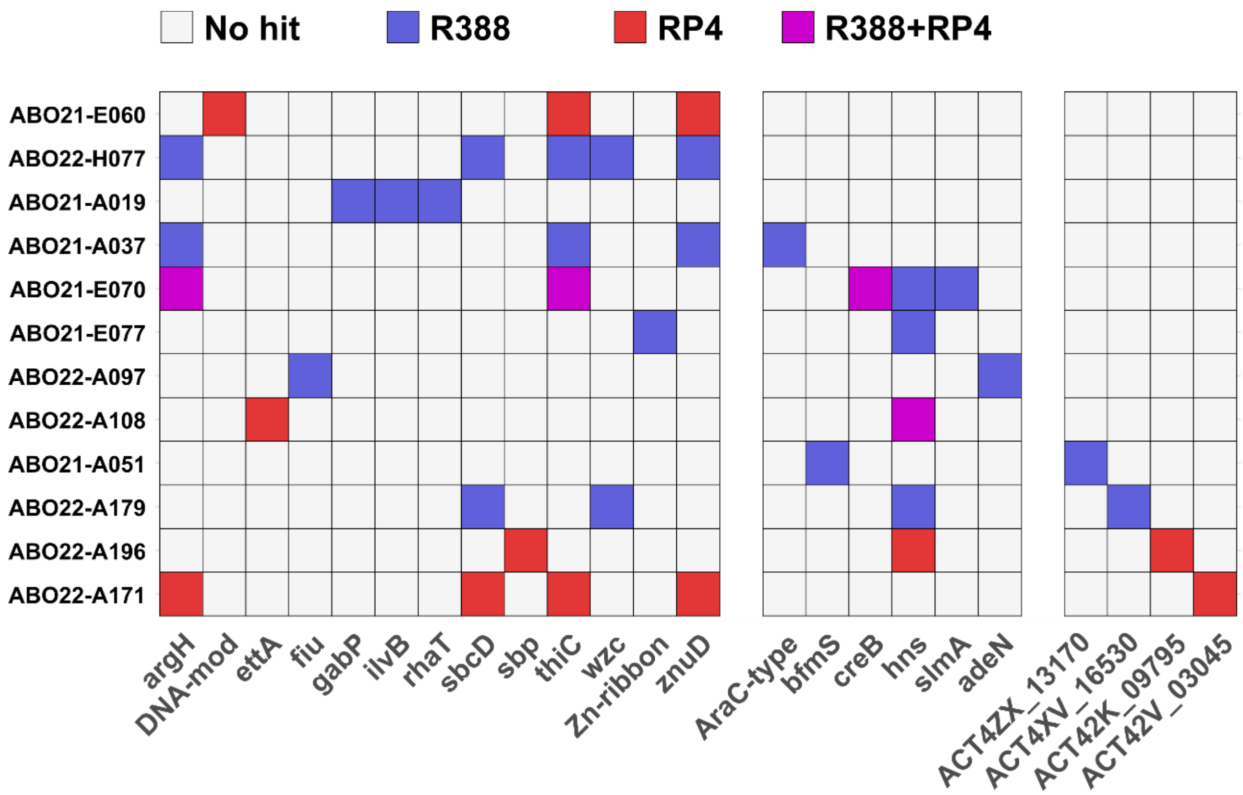


**Supplementary Figure S3.** Exhaustive list of the genes found as regulating plasmid entry using the random mutagenesis approach. On the x-axis are the gene names, on the y-axis are the strains in which they were identified as having an effect. Genes having no effect (white), only regulating R388 (blue) or RP4 (red) entry, or both (pink) are displayed.

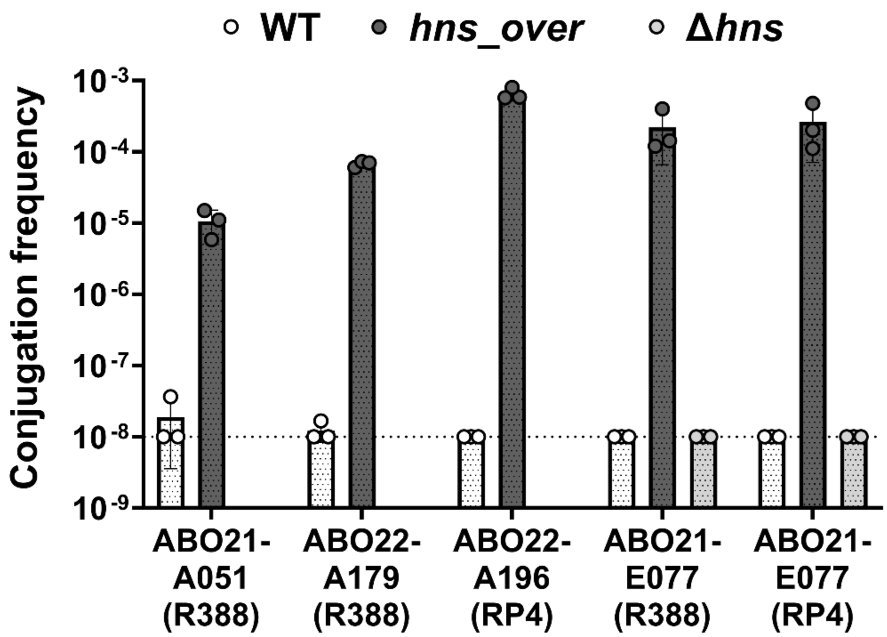


**Supplementary Figure S4.** Impact of *hns* overexpression on conjugation frequency in multiple strains. H-NS overexpression strongly enhances conjugation frequency in various strains (the plasmid for which the transfer efficiency varies is noted below the strain name). This was also performed in the original strains bearing *nmt* (ABO22-A051), *atr* (ABO22-A179) and *ist1* (ABO22-A196). Overexpression in the original strain carrying *ist2* (ABO22-A171) was not possible due to the strain not being naturally transformable. Deletion of *hns* in the ABO21-E077 *hns* overexpression mutant restores the phenotype to levels comparable to the WT strain, confirming the role of this gene in regulating plasmid entry.

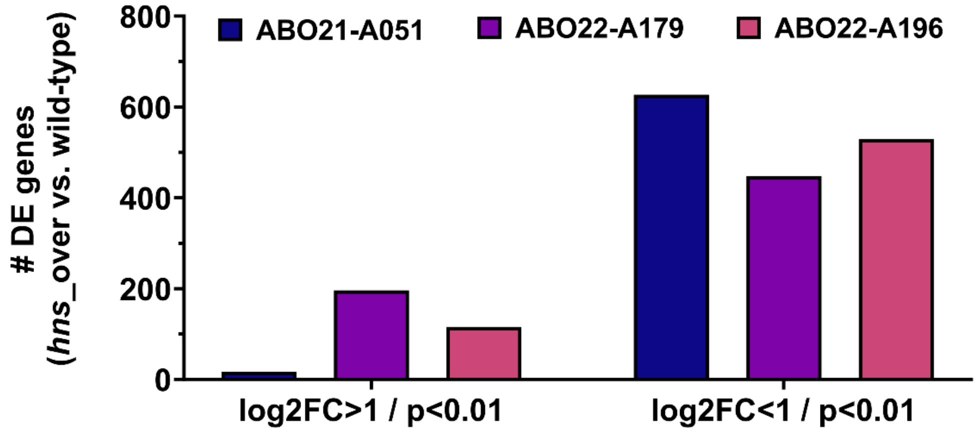


**Supplementary Figure S5.** Number of differentially expressed genes after analysis of the transcriptomic profile of strains in which *hns* overexpression was found to restore plasmid conjugation. Raw RNA-Sequencing data are accessible through accession number PRJNA1456853. Output of the curare pipeline, used to analyze this data, can be found in **Table S2**. Only the genes with |log2FC|>1 and p<0.01 were represented here.

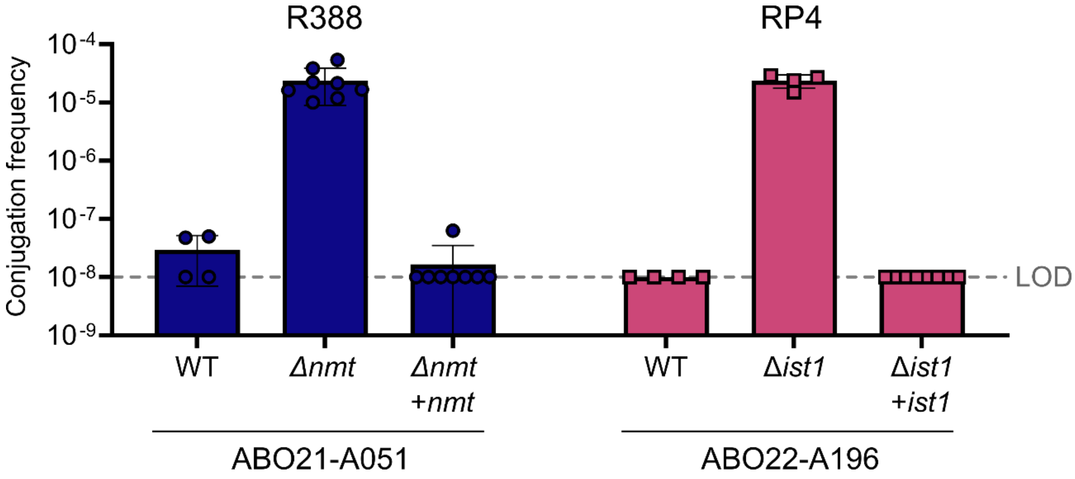


**Supplementary Figure S6.** Confirmation of the role of *nmt* and *ist1* as barriers to conjugation, in their original strains through deletion and complementation. Deletion of the defense genes suppresses their regulatory action on plasmid entry through conjugation, and complementation restores a WT phenotype in the original strains carrying *nmt* and *ist1*. LOD, limit of detection.

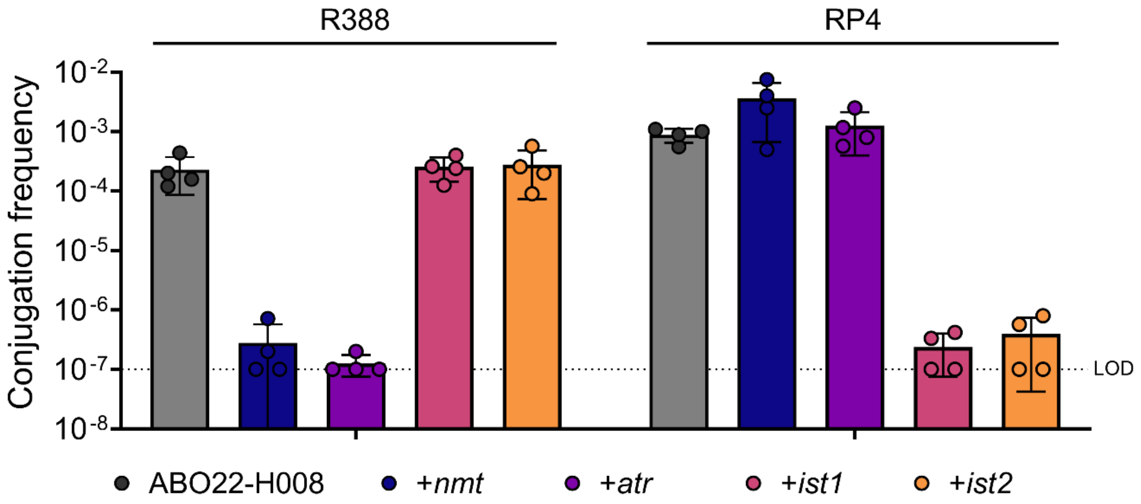


**Supplementary Figure S7.** Confirmation of the role of the four defenses on conjugation, through heterologous expression in the model strain ABO22-H008. LDO, limit of detection.

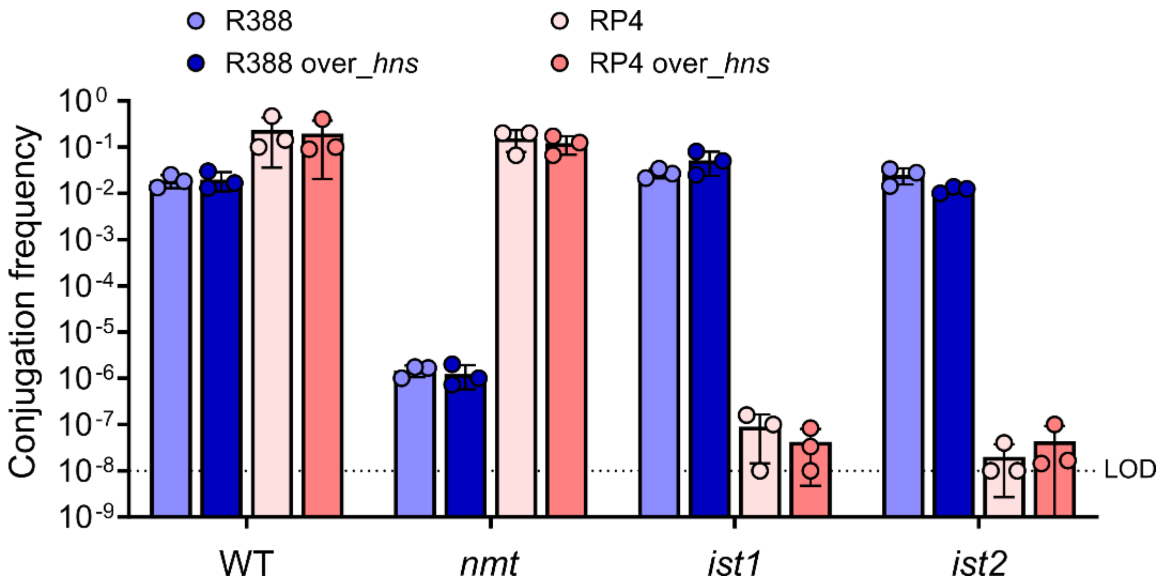


**Supplementary Figure S8.** Effect of H-NS overexpression on conjugation in strain ABO22-H026 expressing the four conjugation defenses. WT, ABO22-H026. LOD, limit of detection.

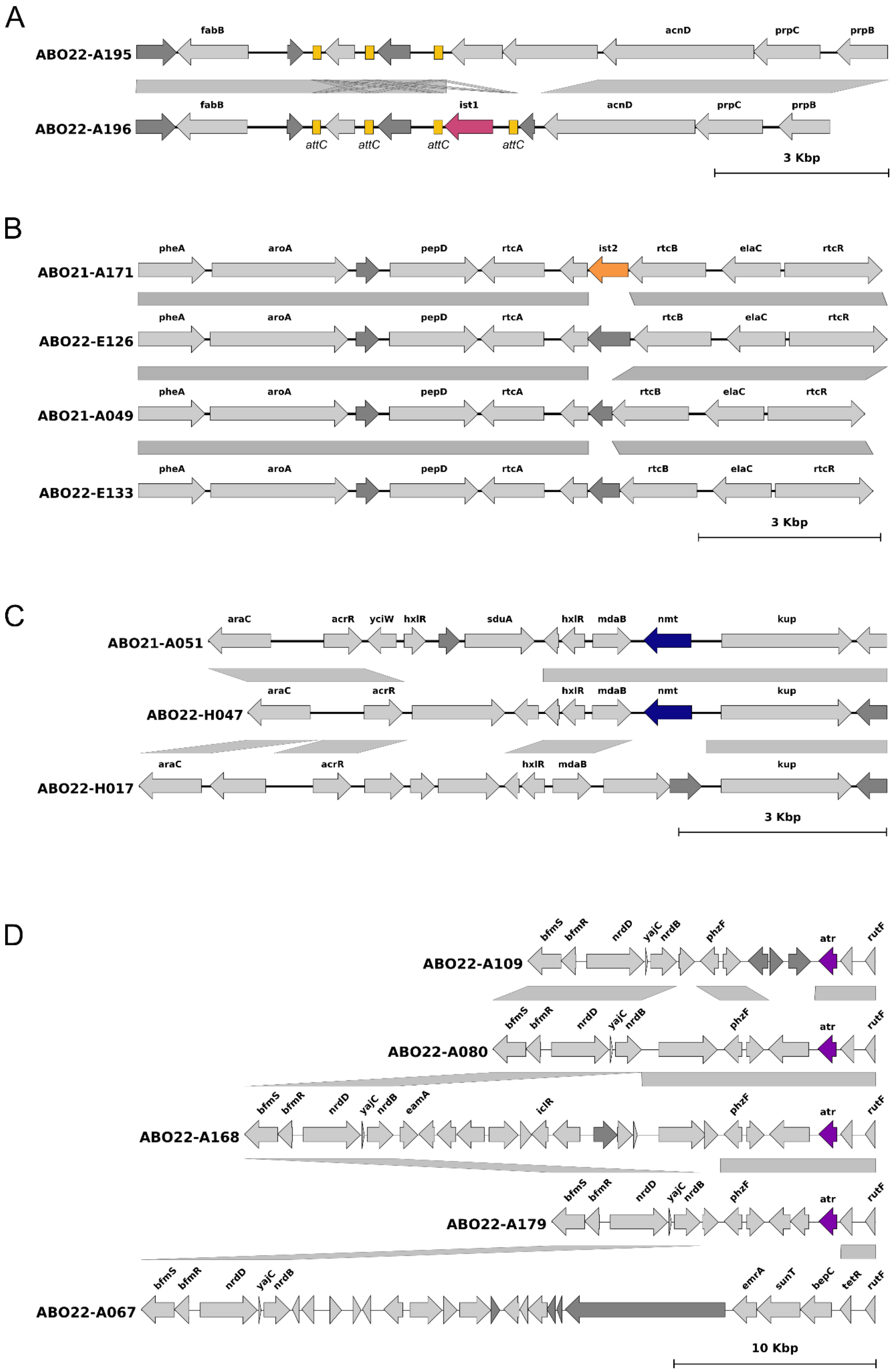


**Supplementary Figure S9.** Genetic environment of the four anti-conjugation defenses. Genetic environment of *ist1* (A), *ist2* (B), *nmt* (C) and *atr* (D) in the original strains in which they were identified and in most closely related strains lacking the gene. Light grey arrows represent annotated genes; dark grey arrows represent genes of unknown function. Yellow boxes represent integron-like *attC* sequences.

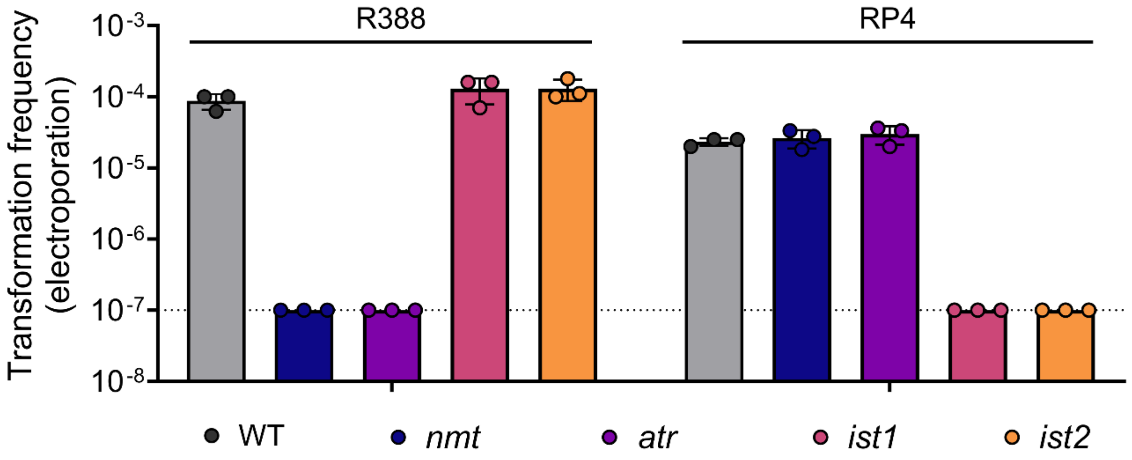


**Supplementary Figure S10.** The four defenses are active against conjugative plasmids delivered by electroporation. Plasmids R388 and RP4 were electroporated in wild-type ABO22-H008 (WT) or expressing the *ist* or *nmt* genes. LOD, limit of detection.

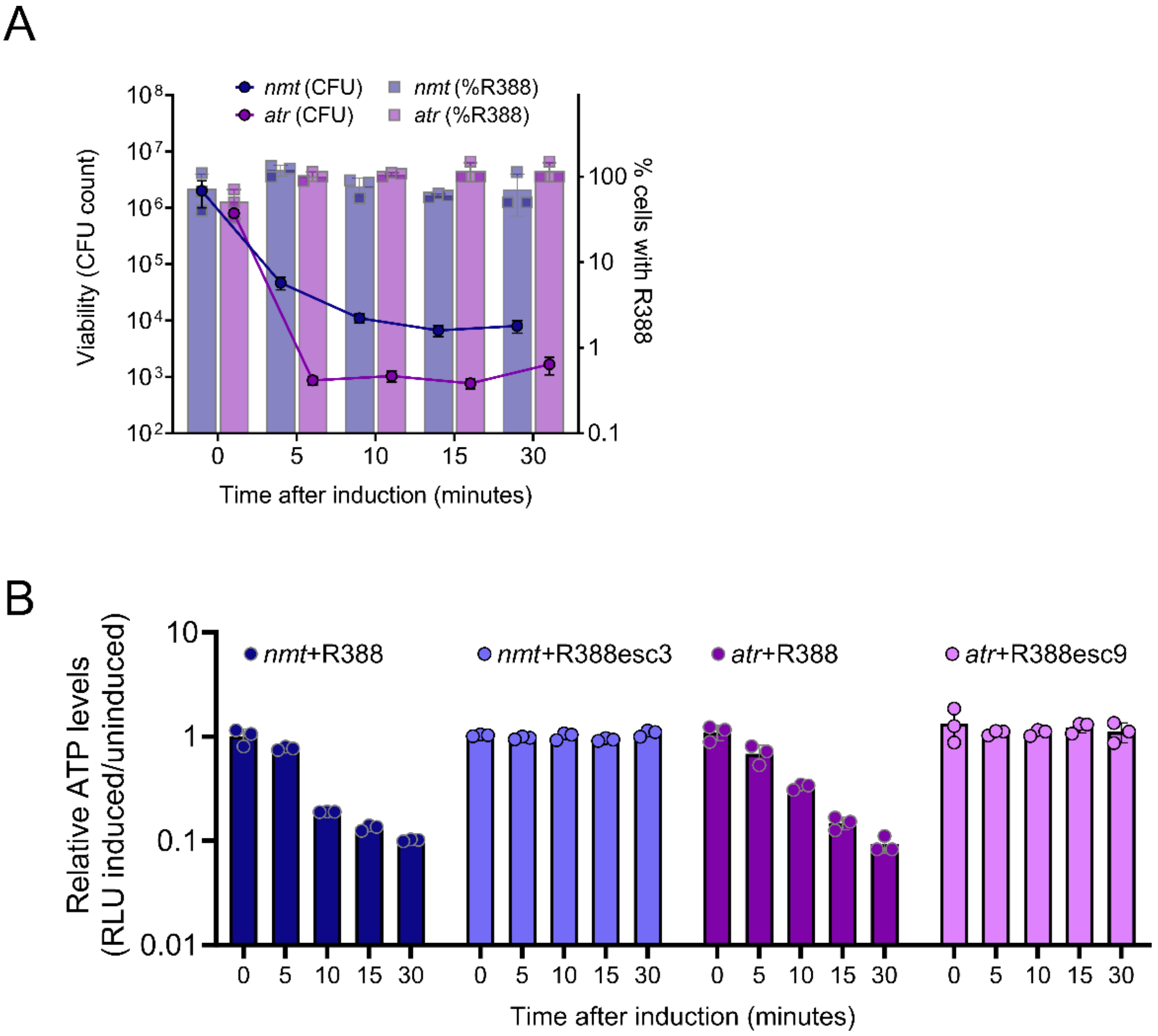


**Supplementary Figure S11.** Namtar defenses cause *virB8*- and *virB2*-dependent cell growth arrest and ATP depletion.

(A). Survival of strains carrying R388 following induction of *nmt* or *atr* genes. Viability is determined by CFU counts, and the percentage of plasmid-bearing cells is determined by CFU counts on LB apramycin versus CFU counts on LB.

(B). Total ATP quantification after induction of *nmt* and *atr* in strain ABO22-H026 bearing R388 or its defense-escaping variants (respectively esc3 and esc9). ATP levels in the cell suspension were derived from luminescence measurements from the BacTiter-Glo™ Microbial Cell Viability Assay (Promega), and the metric represented on the y-axis corresponds to the ratio between the induced and uninduced condition.

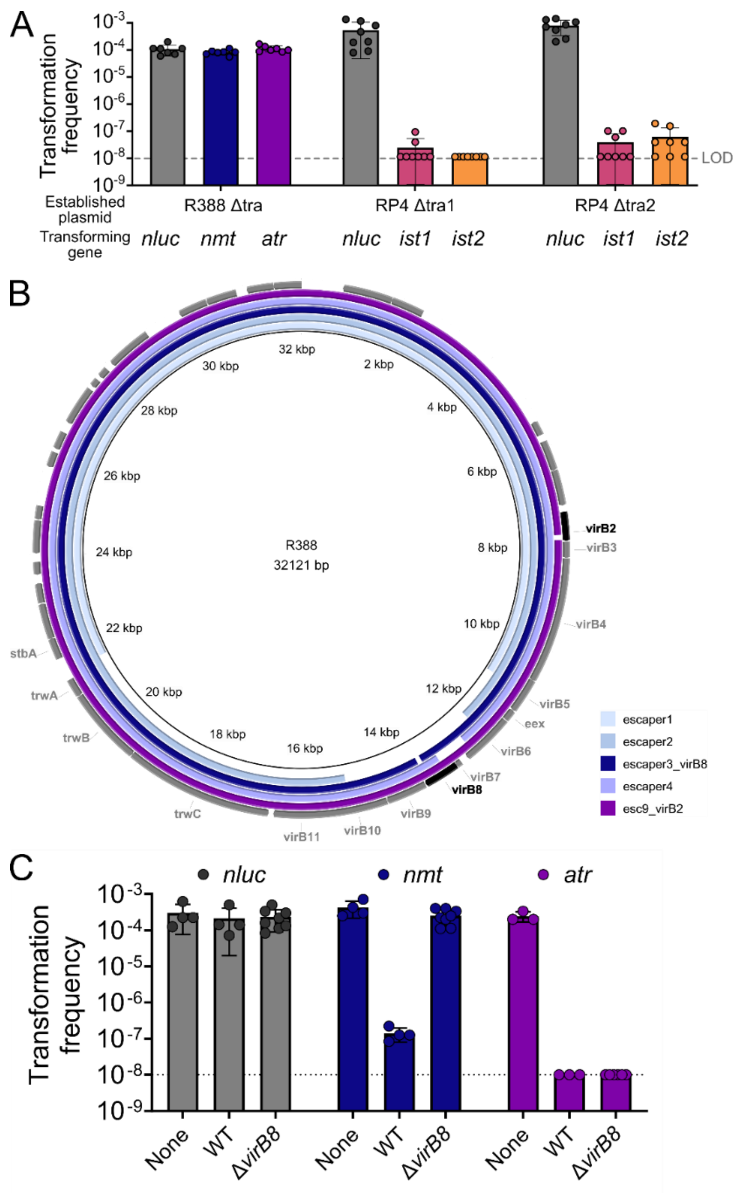


**Supplementary Figure S12.** Deletions in *virB8* and *virB2* allow R388 to escape *nmt* and *atr*, respectively.

(A) Natural transformation of defense-encoding DNA in strains carrying modified versions of R388 or RP4 shows that unlike the anti-RP4 defenses, the anti-R388 defenses target the conjugative machinery. Individual points each represent a single replicate (n=3-5), with the bar end depicting average transformation frequency and SD represented with error bars.

(B) Map of R388 variants escaping defense by *nmt* and *atr* highlights the respective need for *virB2* and *virB8* for defense activation. Variants "esc3" (8-bp deletion in *virB8*, in blue, escaping *nmt*) and "esc9" (118-bp deletion in *virB2*, in purple, escaping *atr*) are represented in color. Escapers 1, 2 and 4, all allowing coexistence with *nmt*, display deletions of varied sizes all including at least a portion of *virB8*.

(C) Efficiency of natural transformation of the *nluc* (control), *nmt* and *atr* genes in strain carrying R388 with a complete deletion of the *virB8* gene.

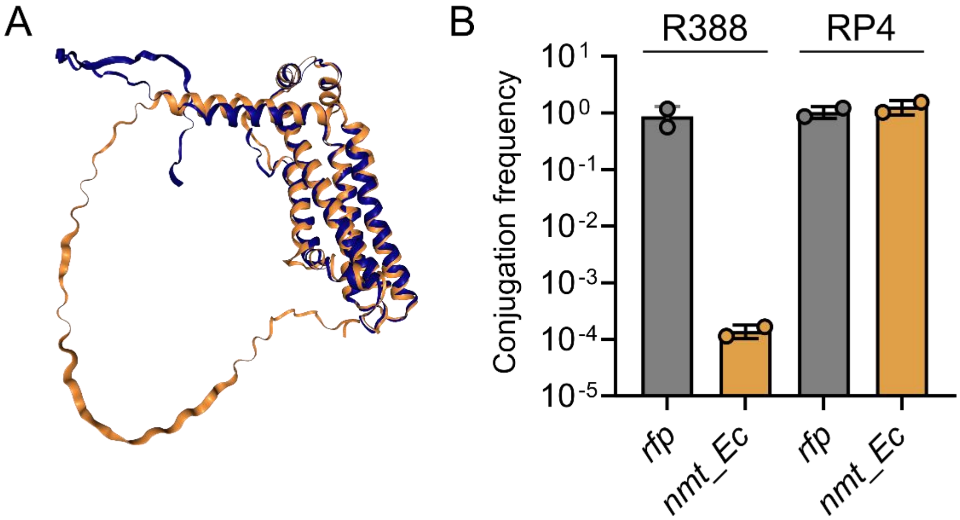


**Supplementary Figure S13. Structural and functional similarities between Namtar homologs in *A. baumannii* and *E. coli*.**

(A) Structures were predicted using AlphaFold 3 and compared using the RCSB Pairwise Structure Alignment tool, which was also used to generate the 3D structure displays. Alignment generated the following stats: RMSD = 1.78; TM-score = 0.75; 27% sequence identity (193/227 *nmt* residues aligned to the *E. coli* homolog).

(B) Conjugation efficiencies of R388 and RP4 in *E. coli* BW25113ΔRM expressing the RFP-encoding gene or the Namtar homolog *nmt_Ec*. Conjugation was performed for 2 hours at 37°C.
